# Linking ringed seal foraging behaviour to environmental variability

**DOI:** 10.1101/2024.09.09.611181

**Authors:** Milaja Nykänen, Marja Niemi, Vincent Biard, Matt I. D. Carter, Enrico Pirotta, Mervi Kunnasranta

## Abstract

**Background:** Foraging rates directly influence animals’ energetic intake and expenditure and are thus linked to body condition and the ability to survive and reproduce. Further, understanding the underlying processes driving a species’ behaviour and habitat use is important as changes in behaviour could result from changes in environmental conditions.

**Methods:** In this study, the dives of Saimaa ringed seals (*Pusa hispida saimensis*) were classified for the first time using hidden Markov models and telemetry data collected on individual dives, and the behavioural states of the diving seals were estimated. In addition, we used generalized additive mixed models on the foraging probability of the seals to identify environmental and temporal drivers of foraging behaviour.

**Results:** We inferred three (in winter) or four (in summer) different dive types: sleeping/resting dives, shallow inactive dives, transiting dives and foraging dives, based on differences in dive metrics logged by or derived from data from telemetry tags. Long and relatively deep sleeping/resting dives were missing entirely in the winter, compensated by an increased proportion of time used for haul-out. We found profound differences in the behaviour of Saimaa ringed seals during the summer open water season compared to the ice-covered winter, with the greatest proportion of time allocated to foraging during the summer months (36%) and the lowest proportion in the winter (21%). The seals’ foraging probability peaked in summer (July) and was highest during the daytime during both summer and winter months. Moreover, foraging probability was highest at depths of 7-30 m in the winter and at depths >15 m in the summer. We also found some evidence of sex-specific foraging strategies that are adapted seasonally, with females preferring more sheltered water areas during winter.

**Conclusions:** We suggest that the foraging behaviour of Saimaa ringed seals is largely influenced by diel vertical movements and availability of fish, and that the seals optimize their energy acquisition while conserving energy, especially during the cold winter months. Further, the seals display some flexibility in foraging strategies, a feature that may help this endangered subspecies to cope with ongoing anthropogenic climate change.

## Background

Foraging encompasses a suite of activities undertaken by animals to locate, acquire, and consume resources essential for maintenance, growth and reproduction. Foraging rates directly influence energetic intake and expenditure and are thus linked to body condition and the ability to survive and reproduce (1). However, foraging effort, success and efficiency are intricately connected to environmental conditions, including temperature and primary productivity (2), which drive prey availability (3–5). Observing foraging in aquatic animals is inherently problematic since this behaviour occurs mostly underwater. For decades, the only source of information on foraging came from direct observations at the water surface, faecal samples, and stomach samples obtained from stranded animals or through commercial harvests. However, recent developments in biologging and animal telemetry have opened new possibilities for studying movement and foraging patterns of aquatic animals (6). Movements of individuals can be monitored by deploying tags that collect information on location, activity (*e.g*., dives), acceleration or the surrounding environment (7,8). Subsequently, it is possible to infer an individual’s behavioural state by assuming that different behaviours are associated with specific characteristics of its movement patterns. For example, when analysing location data, travelling is typically characterised by directed movement trajectories, whereas convoluted trajectories are associated with foraging activity on patchy food resources (9). With the accumulating evidence for global biological impacts of climate change (10), predicting future responses of plant and animal populations to ongoing environmental change is increasingly important. Climate change is especially affecting arctic and subarctic mammals such as ringed seals (*Pusa hispida*) that rely on ice platforms and snow drifts for successful pupping, nursing, resting and moulting (11,12). The endemic lacustrine Saimaa ringed seal (*P. h. saimensis*) has suffered dramatic declines over the past century from a long-term effective population size (prior human impacts) estimated at 1270 seals (95% CI: 220–8850, (13)), although the population is currently increasing due to successful conservation efforts (14). The Saimaa ringed seal is currently listed as Endangered (International Union for Conservation of Nature Red List), with a population estimate of ~480 individuals (15). The population is increasing by ~3% annually (16) but at a lower rate than the potential annual growth rate of seal populations (17) due to ongoing anthropogenic threats, such as bycatch, especially in recreational gillnet fishery, fragmentation of suitable habitat and climate change (16). To improve the survival of juveniles that are especially affected by gillnet bycatch mortality, various fishing restrictions are implemented in Lake Saimaa, including an annual seasonal ban on gillnet fishing lasting from the 15th of April until the 30th of June, a period thought to be the most critical for juvenile dispersal. Despite the restrictions, estimated annual juvenile bycatch mortality remains at 10-13% (18).

Saimaa ringed seals display a high degree of site fidelity (19,20). They have stable home ranges of around 90 km^2^, with 5 km^2^ core areas, during the open water season but may also undertake sporadic trips lasting for a few days outside their normal range (21). In winter, when the lake is (normally) frozen over, the seals use snow lairs for resting and pupping (16). During this time, their movements are more restricted, with winter home ranges less than a tenth, and core areas less than half the size compared to the open water season (22). In the spring, the seals undergo an annual moult, during which they spend extended periods of time hauled out on land after the ice breaks up (23). After the moult, the seals are assumed to focus their activity on foraging, with prey consumption being highest in autumn (24). Saimaa ringed seals are opportunistic feeders, and, according to analyses of stable isotopes and digestive tract contents, they feed exclusively on small schooling fish (24,25).

Foraging of ringed seals has previously been inferred from dive patterns, with convoluted dives hypothesised to involve local search within prey patches, pursuit of prey, and/or agonistic interactions with conspecifics (26). Also Saimaa ringed seals’ dives have been associated with different types of behaviour based on dive depth, duration and shape of the dive profile, with three different behavioural categories inferred; travelling, foraging, and resting (27). However, these classifications have been based on manual processing of dives and thus limited to a few hundred dives collected from three to five animals. Recent advances in analyzing Global Positioning System (GPS) tagging data (28–31) have opened new possibilities to look for patterns in movement data and identify foraging and other behavioural states. Hidden state, or hidden Markov, models (HMMs) can be used to identify discrete movement patterns in data (32,33) and thus allow ecologists to infer and classify behavioural states from large datasets containing many thousands of observations. In a comparison between different movement modelling approaches, ground-truthed against direct metrics of foraging activity, HMMs have performed the best in estimating the behaviour of diving seabirds (34). A key advantage of these models over other analytical options is the ability to compute the transition probabilities between different behavioural states as a function of covariates, thus enabling the assessment of the role of environmental variables driving the decision-making process of animals (31,35). Previous studies on the movement ecology of Saimaa ringed seals have concentrated on their distribution, home range size, foraging habitat range and effects of disturbance (*e.g*., 21,36). However, the behavioural patterns during dives of ringed seals inferred from movement data have yet to be quantified in the context of varying temporal and environmental conditions. HMMs that utilize vertical dive data (37,38) may be more suitable for a species such as the Saimaa ringed seal whose movement paths are unlikely to contain clear travelling segments due to restricted habitat and small home ranges, and whose area-restricted movements may encapsulate multiple behaviours (*e.g*., foraging, resting), making it hard to quantify foraging rates. The use of dive data can elucidate this.

Understanding the underlying processes driving a species’ habitat use is important as changes in behaviour could result from changes in environmental conditions. For example, environmental change has been linked to changes in foraging phenology (39). As top predators, ringed seals can act as sentinels of climate change due to their sensitivity to environmental variability (40), which is exacerbated by their dependency on ice and snow. Moreover, understanding the temporal and environmental drivers of behaviour will help focusing conservation measures put in place to protect this endangered subspecies. Therefore, the aim of this study was to uncover the underlying temporal, spatial and environmental factors that influence foraging of ringed seals. We focused on two distinctive periods of the seal’s annual cycle; the summer open water season, when the seals are thought to forage extensively, replenishing their lipid reserves after the annual moult; and the ice-covered winter period, during which the seals breed.

## Methods

### Study area

Lake Saimaa (Fig. 1), the largest lake in Finland with a surface area of 4,400 km^2^, is approximately 180 km long and 140 km wide, with a mean depth of 12 m and maximum depth of 85 m. The lake is extremely fragmented, enclosing almost 14,000 islands and consisting of several basins interconnected by narrow straits (41). There are six towns located around the lake, with a combined human population of over 286,000, and nearly 70,000 vacation homes (42) are scattered along the shorelines and islands. Lake Saimaa typically freezes over in December - January (see Table S1, Supporting information, for the dates in this study), and the ice breaks up and melts in April - May. Like many boreal lakes, Lake Saimaa is dimictic, with complete overturns of the water column occurring in the spring and autumn. During these overturns, water is mixed, and nutrients are released from the deeper layers to the surface, which affects the phyto- and zooplankton species assemblages (43,44), and, in turn, higher trophic levels (45).

**Fig. 1.**
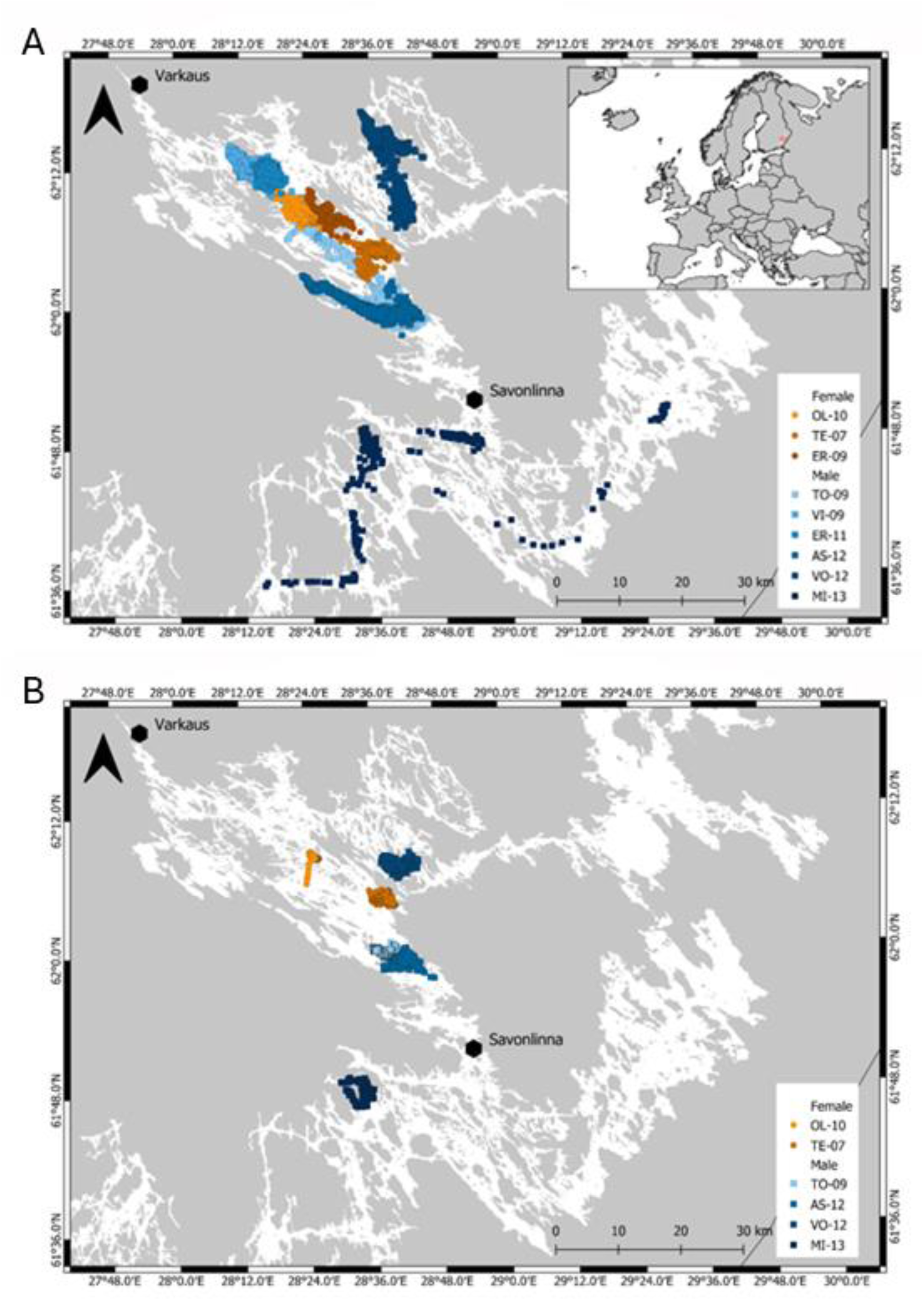
Locations of inferred foraging dives of Saimaa ringed seals in (A) summer and (B) winter. Gray area represents land and white area water. Towns of over 20,000 inhabitants are indicated on the map.

### Data collection

Fastloc GPS-GSM (Global System for Mobile communications) phone tags (Sea Mammal Research Unit Instrumentation, St Andrews, UK) were deployed on nine adult ringed seals (six males and three females) captured in central Lake Saimaa soon after their annual moult in late May - early June 2009-2013 (Fig. 1). The seals were captured and handled for the deployment in accordance with the Finnish environmental authorities, Centre for Economic Development, Transport and the Environment (ESAELY/433/07.01/2012 and ESA-2008-L-519-254), and the Project Authorisation Board (ESAVI/8269/04.10.07/2013 and ESAVI-2010-08380/Ym-23). The tags were attached by two-component epoxy glue to the dorsal pelage, with the antennae pointing backwards (except for one individual, OL10, Table S1, Supporting information) due to the potential damage caused by ice-cover in the winter. After tagging, seals were released in the vicinity of their capture sites. The tags collected data over 136-328 days (Table S1, Supporting information) dropping off with the old fur during the next moult at the latest.

The phone tags were equipped with a GPS receiver and wet-dry and pressure sensors, providing geo-referenced summaries of individual dives and haul-out events via the GSM phone network (46). The programming of the tags (GPS recording and GSM call interval) varied slightly between years due to tag development (see Table S1, Supporting information). Furthermore, due to the seals’ diving and variation in satellite availability, GPS positions were recorded irregularly. On average, 19 locations per day in summer and 5 locations per day in winter were recorded (Table S1, Supporting information). When the tag’s wet-dry sensor determined that the animal was submerged, the pressure sensor also recorded depth. Dive start times were defined as when the wet-dry sensor was wet continuously for 8 s and the depth was below a threshold of 1.5 m, and end times as when the pressure sensor recorded depth above 1.5 m. Each dive was summarised using depth bins at nine equally spaced time points throughout the dive, and maximum diving depth, duration, and time-depth summary were transmitted through the GSM network.

### Data processing and retrieval of environmental covariates

The seals’ dive locations (start and end) were estimated by linearly interpolating the GPS positions using the manufacturer’s software. However, due to the complex lake landscape some of the dive locations were estimated on land. These locations were moved off-land using the package ‘pathroutr’ (47). Dives occurring within the first 24 h after the deployment were discarded to avoid including any behaviour potentially impacted by the tagging procedure. Due to the presence of unusually long dives, we filtered the dives for surfacing events that were potentially missed by the tag, *sensu* (48). These potentially incorrect dive records were defined as the ones during which the animal stayed within the dive threshold depth (1.6 m) throughout the dive, or when the dive duration exceeded 400 s and at least one depth reading (excluding the first descending and last ascending phases) in the upper 25% of the dive occurred without surfacing. This filtering removed between 0.06 and 2.58% of an individual’s dives in the summer and between 0.79 and 16.43% of dives in the winter, depending on the animal.

After filtering the dives, we retrieved a range of weather variables (air temperature (°C), rain accumulation (mm), and wind speed (m/s) and direction) from the open portal of the Finnish Meteorological Institute (http://opendata.fmi.fi/wfs). A custom R-script was used to extract the data from all weather stations located within 50 km of the corrected dive start locations to the nearest hour. The extracted values (*v*) were then interpolated for each variable at each location using averaging with square-exponential weighting by distance: 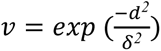, where *δ* is the bandwidth of 50,000 m and *d* is the distance (in m) from the dive location to the weather station. We also retrieved the nearest water depth value (m) for each dive location using the ‘sf’ R-package (49) from the depth points (Sounding_P) raster layers corresponding to the areas of Lake Saimaa, downloaded from the open data portal (https://julkinen.traficom.fi/oskari/?lang=en) of Finnish Transport and Communications Agency Traficom. In addition, we derived an index of “openness” of the water area for each 100 m ×100 m cell of Lake Saimaa raster using the focal() function from ‘raster’ R-package (50), and extracted the values for each corrected dive location using extract() from ‘terra’ package (51). Low values of this variable indicate sheltered areas surrounded by land, while high values indicate open areas.

### Statistical modelling

We used a two-step modelling approach to investigate the role of temporal and environmental drivers on the diving behaviour of Saimaa ringed seals. First, we fitted HMMs to classify diving behaviour. We included temporal (Julian day and hour of the day) and the environmental variables as covariates in the HMMs affecting the state transition probabilities to investigate whether they could act as cues for the seals to initiate different behaviours, especially foraging activity. Second, we fitted generalized additive mixed models, GAMMs (52), on the foraging probability of seals and including the same covariates as in the HMMs, plus a factor covariate sex, while treating each seal ID as a cluster of observations. By fitting a GAMM *post hoc*, we were able to account for the effect of sex and for random variation among animals, permitting us to make population-level inferences regarding the drivers of foraging behaviour of Saimaa ringed seals.

#### Classifying behaviour with hidden Markov models

We fitted a HMM using pooled dive data collected from nine seals (three females and six males) during the summer before the lake autumn turnover (June–September). In addition, a separate HMM was fitted to the pooled data collected from six of the seals (two females and four males) during the ice-covered period in the winter (with the temporal extent varying across years, see Table S1, Supporting information). Two separate models were run due to the differences observed in plots of summary statistics of the dive parameters (dive duration (s), and maximum dive depth (m)) logged by the GPS-GSM tags, indicating a change in dive behaviour in winter compared to summer. We ran both HMMs with three different data streams: dive duration, maximum dive depth, and dive sinuosity (or “wiggliness”), which was defined as the vertical distance (m) covered at the bottom phase of each dive and calculated as per (37,53). Dive sinuosity is considered a good proxy for prey chasing behaviour (37). We selected a gamma distribution for the data streams as they were all non-negative continuous values. To overcome numerical problems of the HMM fitting algorithm and to minimize residual autocorrelation (visible in the autocorrelation function plots for pseudo-residuals in preliminary models run at the scale of individual dives), we divided the dives into batches of ten for the summer model (31,54). Also, if the time between consecutive dives exceeded one hour, they were placed into separate batches. For the winter model, we divided the dives into batches of five as this produced a sufficient reduction in residual autocorrelation. The data streams and the environmental covariates were summarized at the batch scale by averaging them across the dives in each batch, except for the wind direction for which the median was taken to capture the prevailing direction. Step length or turning angle were not used as data streams due to the gaps and error in GPS locations and the complexity of the lake landscape, with numerous narrow straits and islands forcing the seals to move extremely directionally, ultimately limiting the usefulness of these movement metrics for describing separate states. Based on a previous study (27), we assumed that the model would be able to classify at least three different latent states that differed in their dive metrics and that were inferred to represent different dive behaviours: sleeping/resting, transiting and foraging. Sleep/rest dives were assumed to be the deepest and longest with very few vertical movements occurring during the bottom phase of the dive (low sinuosity). Foraging dives were also assumed to be long and relatively deep but, in contrast with sleep/rest dives, they were assumed to have high sinuosity values. Transit dives were assumed to be of intermediate duration, depth and sinuosity. In addition, in preliminary models run on a reduced dataset, an additional latent state was identified. The dives in this state, named hereafter as ‘shallow inactive’ did not match the expected characteristics of the above dive categories but were characterized by shallow depth, short dive duration and low dive sinuosity. We ran the models with constraints *sensu* (55,56), by constraining the gamma rate mean of dive sinuosity to be lowest for sleep/rest dives and highest for foraging dives (μ_sleep_ < μ_shallow inactive_ < μ_transit_ < μ_forage_), and constraining the gamma rate mean of dive duration as μ_transit_ < μ_forage_. The initial values for the model parameters were selected based on estimates from preliminary models run on data from two of the seals (data from individual dives in June). Specifically, we used estimates from the model with the lowest Akaike’s Information Criterion (AIC; (57)) score among 50 iterations with randomly selected initial values drawn from a uniform distribution with defined lower and upper bounds (following (28)), with the exception of dive sinuosity, for which the initial values were set based on the above constraints and averages calculated from sequences of individual dives.

In addition to running a model without any covariates (the null model), we included the standardized, batch-averaged water depth, air temperature and openness index, and B-splines of hour of the day and Julian day (or cumulative day in the winter model, i.e., we did not reset the day to 1 on January 1st in order to have a continuous time series), in a stepwise process starting from the simplest model with only one covariate (air temperature) and adding terms one by one. The relevance of the covariates was judged from the plotted transition probabilities and using AIC with lower values indicating a better fitting model. Covariates were kept in the model if their inclusion reduced the AIC-value by at least two units (58). Following preliminary analysis, we excluded rain accumulation and wind speed and direction from the models due to the large number of missing values (altogether 76% and 95% missing values in summer and winter, respectively). We ran the models with 2-4 possible states and selected the number of states best representing the data by visually examining the overlap between state-dependent distributions. We did not use the AIC to select the number of states, because standard information theoretic criteria generally prefer models with a larger number of states that are difficult to interpret biologically (59,60). The Viterbi algorithm was then used to assign the most likely sequence of batch states for each deployment (61).

#### Effect of temporal and environmental covariates on foraging (GAMMs)

The latent state estimated at each dive batch was converted to a binary response variable of putative foraging vs non-foraging (1 or 0), which was then modelled as a function of sex and a set of temporal and environmental covariates hypothesized from the literature as potential drivers of foraging behaviour. Specifically, we included the same covariates investigated in the HMM analysis, and fitted the GAMMs in a binomial framework using the bam() function from package ‘mgcv’ for R (52,62), where each individual Saimaa ringed seal was included as a random intercept. Because preliminary analysis suggested some temporal autocorrelation in model residuals (summer ρ = 0.496, winter ρ = 0.454), a first-order autoregressive correlation structure (AR1) was fitted within each level of the random effect (i.e., each seal ID). Since the response variable in our model was binary (*i.e*., non-Gaussian), the AR model is applied to the working residuals and corresponds to a Generalized Estimating Equations (GEE) approximation (52,63). We assessed the possible correlation between model covariates with the ‘concurvity’ function of the ‘mgcv’ package (52,62) and found no evidence of concurvity between any of the variables (with all concurvity estimates of <0.35 in the summer model and <0.50 in the winter model). We used penalised thin-plate regression splines with shrinkage (64) to model the relationship between the binary response and each of the explanatory variables, with the exception of hour, which was fitted using a cyclic spline to capture daily patterns. Separate covariate smooths were fitted for each sex via the ‘by’ argument. The performance of the final model was assessed using a confusion matrix, which compared the occurrence of foraging predicted by the GAMMs with the behavioural state estimated by the HMM. Further, the goodness-of-fit of the model was assessed by calculating the area under the receiver operating characteristic curve (AUC) using the package ‘ROCR’ for R (65). The contribution of each explanatory variable was visualised using partial residual plots generated using ‘mgcv’.

The uncertainty resulting from the state classifications from the HMM (which determined the value of the response variable used in the GAMM) was propagated following (35), that is, using the estimated probabilities associated with each state obtained from the HMM in a multinomial draw for each dive batch and repeating the procedure 100 times. At each draw, new state estimates were returned, which were used to rerun the GAMM. The AUC, confusion matrix and predictions were also calculated at each iteration, and summaries of these metrics across the 100 iterations provided an indication of the effects of state classification uncertainty on the final results. Finally, uncertainty in the GAMM predictions was visualized by overlaying the effect of each model covariate on the predicted foraging probability (mean and 95% prediction interval) from the 100 runs.

#### Foraging area overlap

To investigate if there was evidence of resource partitioning between the seals and to quantify the extent to which the foraging areas of different seals overlapped in the summer vs winter, we calculated a bivariate kernel utilization distribution (UD), with kernel density estimator calculated at 95%, based on the locations associated with foraging dives inferred for each individual. Foraging area overlap was estimated for each possible seal pair (36 pairs in the summer and 15 in the winter) using Bhattacharyya’s affinity (BA; (66)) where 0 equates to no overlap and 1 equates to 100% overlap in the UDs. Further, to compare space use during different seasons, the overlap in foraging areas between summer and winter for each individual seal was estimated using BA. All overlap analyses were done using R-package ‘amt’ (67).

## Results

### Behavioural states

According to AIC scores, the HMM that included Julian day, hour of the day, depth, and openness of the water area was selected for the summer data while excluding air temperature (Table S2, Supporting information). The summer model differentiated four latent states (Fig. 2A-C). The first state, which was interpreted as sleep/rest, included the longest dives and was also characterized by deep maximum depth and very low vertical sinuosity. The second state, interpreted as shallow inactive, included dives of short to intermediate duration, relatively shallow depth and low sinuosity, while the third state, interpreted as transit/other, contained dives of intermediate dive duration, depth and sinuosity. The fourth state was characterized by long sinuous dives occurring at the deepest dive depths (>15 m in the summer) and was interpreted as foraging. The results of the summer model show that Saimaa seals were more likely to remain in the foraging state around late July and tended to switch to foraging from any other state early in the morning and with increasing openness of the water area (Fig. 3; Figs. S1-S2, Fig. S4 in Supporting information). There was a positive nonlinear correlation between water depth and switching to foraging from all the other states except for the sleep/rest state (Fig. S3, Supporting information). In general, the probability of foraging increased with depth up to about 20 m, after which it started to decline (Fig. 3). The seals spent proportionally the least time engaged in sleeping/resting behaviour (9.2%) followed by shallow inactive (18.8%) behaviour. In the summer, the greatest proportion of time was allocated to foraging (36.2%) and transit/other behaviour (35.7%).

**Fig. 2.**
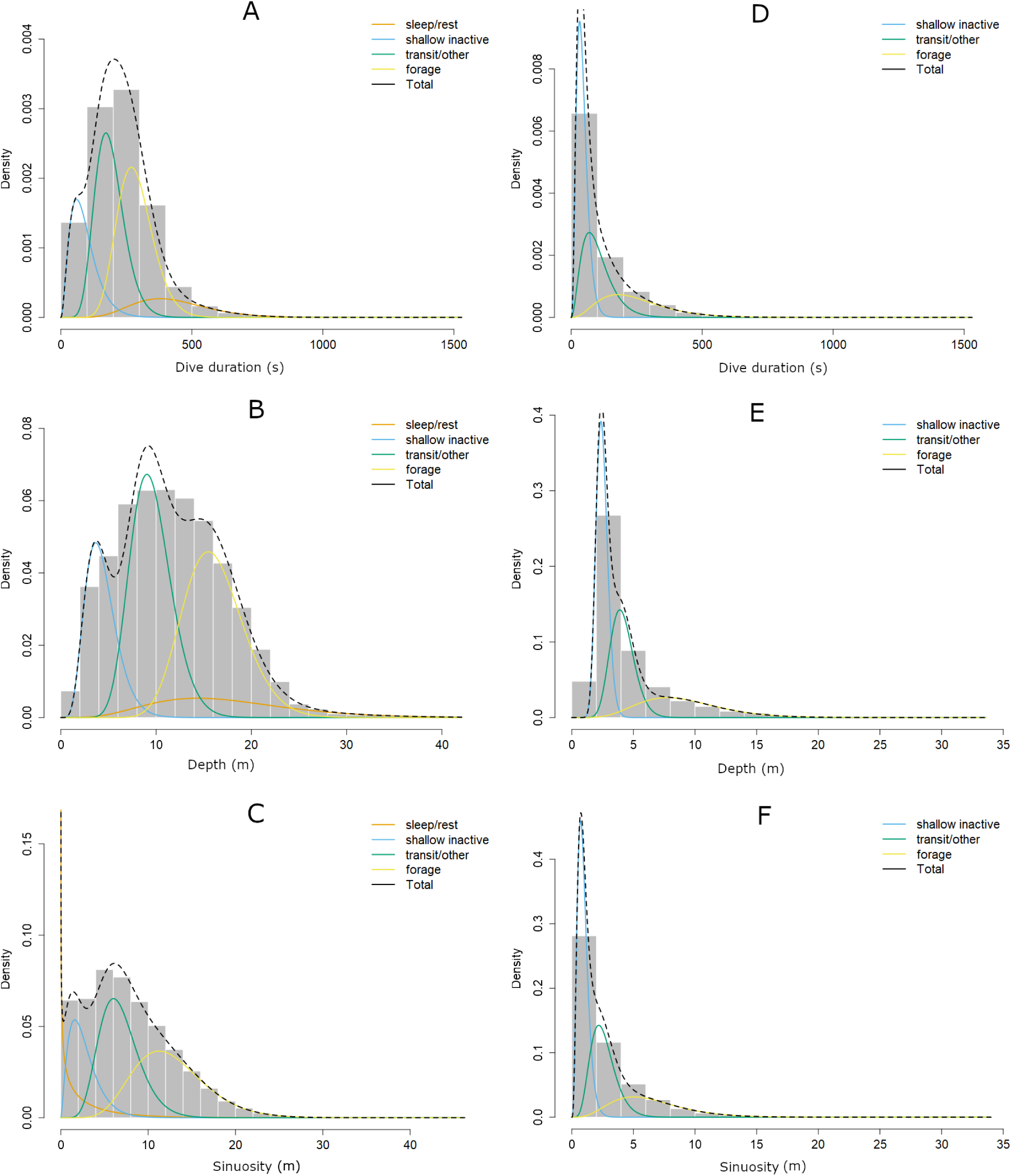
Plots of state-dependent distributions of dive duration, maximum depth and dive sinuosity (summarized as mean for each batch of dives) for Saimaa ringed seals from the hidden Markov model during summer (A – C) and winter (D – F).

**Fig. 3.**
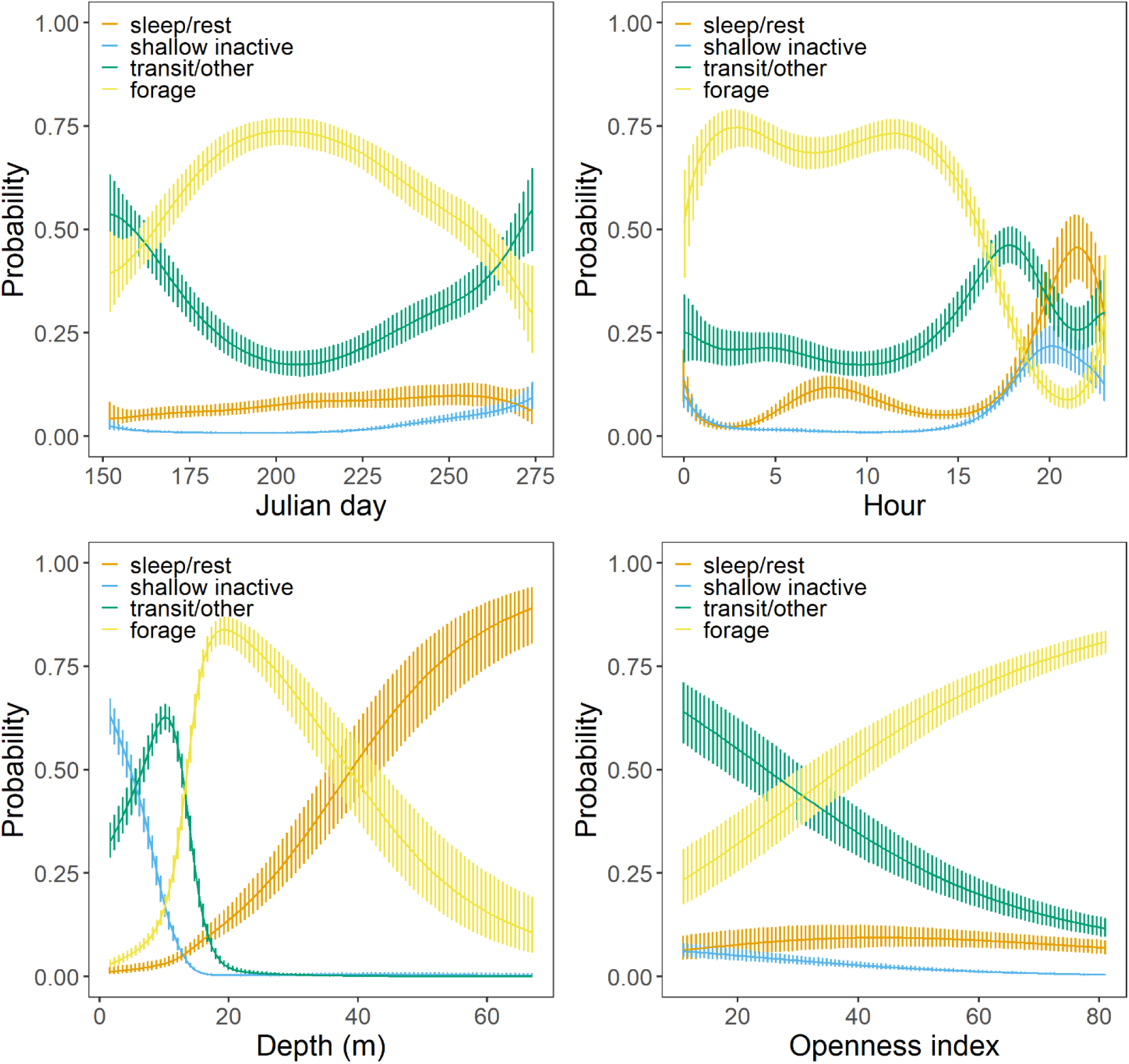
Stationary (long-term) state probabilities of the HMM for Saimaa ringed seals during the summer period, given as functions of the four covariates for state transition probabilities. The vertical lines depict pointwise 95% confidence intervals.

The HMM that included the effect of all temporal and environmental covariates was preferred over simpler models for the winter data based on AIC scores (Table S2, Supporting information). The HMM classified winter dive batches into three different states, interpreted as shallow inactive, transit/other and foraging, with the long and deep sleeping/resting dives typical of the summer period missing entirely (Fig. 2D-F). Moreover, the dives identified as foraging were shorter, shallower and less sinuous compared to the foraging dives occurring in the summer. As opposed to the summer model, there was no clear seasonal pattern associated with any of the states in the winter model (Fig. 4; Fig. S5, Supporting information). However, there was a similar diel pattern to the summer model as the probability of engaging in, or switching from other states to, foraging was highest in the morning (Fig. 4; Fig. S6, Supporting information). There was a slight negative correlation between air temperature and the probability of switching from transit/other state to foraging as well as remaining in foraging (Fig. S7, Supporting information). The likelihood of foraging and switching to foraging increased non-linearly with water depth (Fig. 4; Fig. S8, Supporting information), and there was a positive correlation between the openness of the water area and the probability to switch from transit/other state to foraging as well as remaining in the foraging state (Fig. 4; Fig. S9, Supporting information). The proportion of time spent in different behavioural states was different in the winter compared to the summer, with 20.8% of time allocated to foraging, 33.5% to transit/other and 45.7% to shallow inactive.

**Fig. 4.**
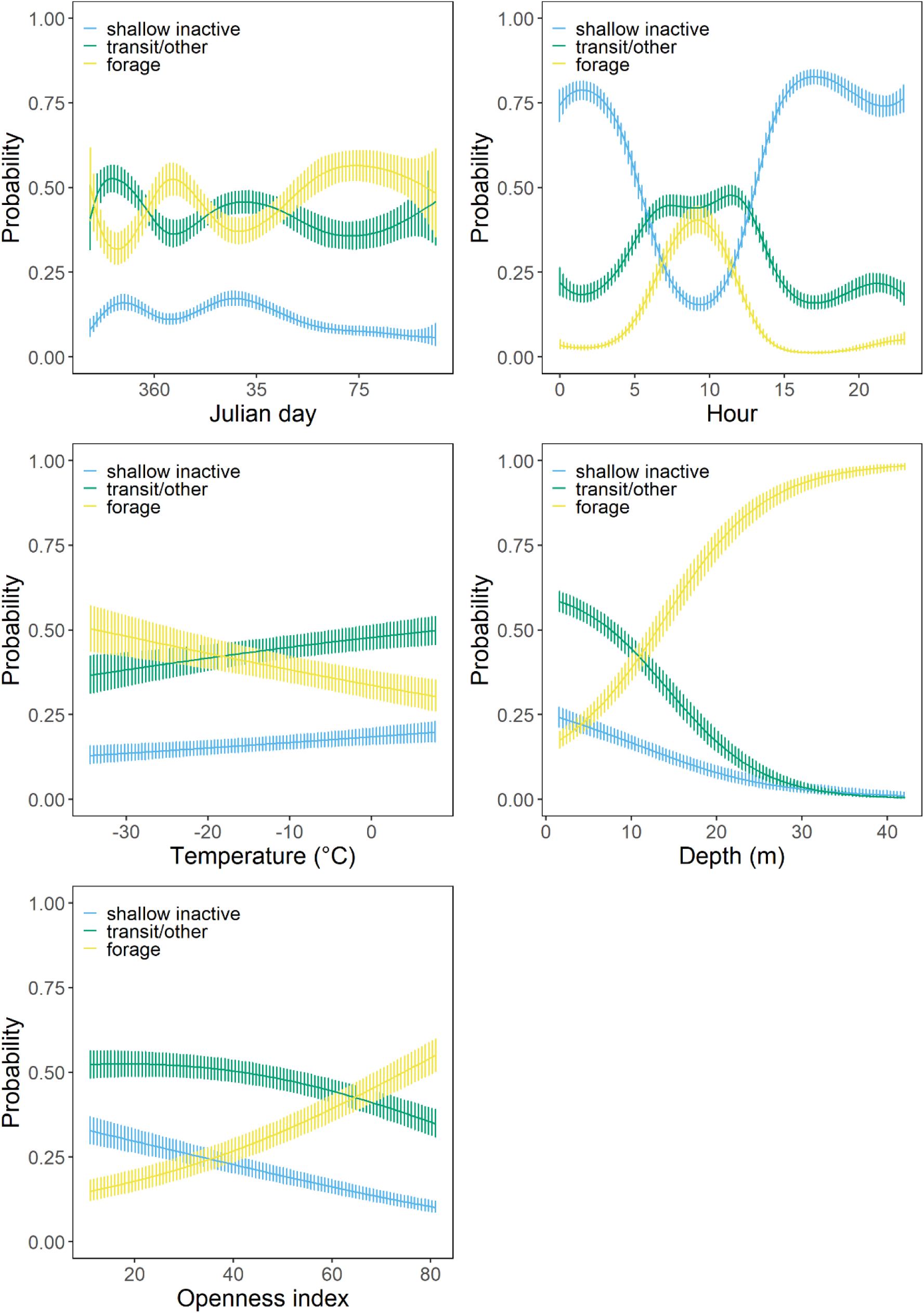
Stationary (long-term) state probabilities of the HMM for Saimaa ringed seals during the winter ice-covered period, given as functions of the five covariates for state transition probabilities. The vertical lines depict pointwise 95% confidence intervals.

### Environmental variables associated with foraging

All temporal (Julian day and hour of the day) and environmental covariates (air temperature, water depth and openness index) interacting with sex were retained in both summer and winter GAMM as none of them were shrunk to zero. In the summer, the seals were more likely to engage in foraging behaviour around Julian day 200, which corresponds to mid to late July, with a slight delay in the peak of foraging probability for males compared to females (Fig. 5A). The cyclic smooth for hour suggested a higher chance of seals engaging in foraging behaviour from the early hours of the morning until mid-afternoon (Fig. 5B). Air temperature had a small but non-zero effect on the foraging probability of males with a marginally significant positive linear relationship (effective degrees of freedom [edf] = 0.71, χ^2^ = 4.33, p = 0.07), whereas the effect was small and non-significant in females (edf = 0.41, χ^2^ = 0.67, p = 0.23) (Fig. 5C). Foraging probability was highest at depths >15 m for both females and males (Fig. 5D). Female seals were more likely to engage in foraging behaviour in more open water areas, while in males foraging probability peaked in both sheltered and open areas (Fig. 5E).

**Fig. 5.**
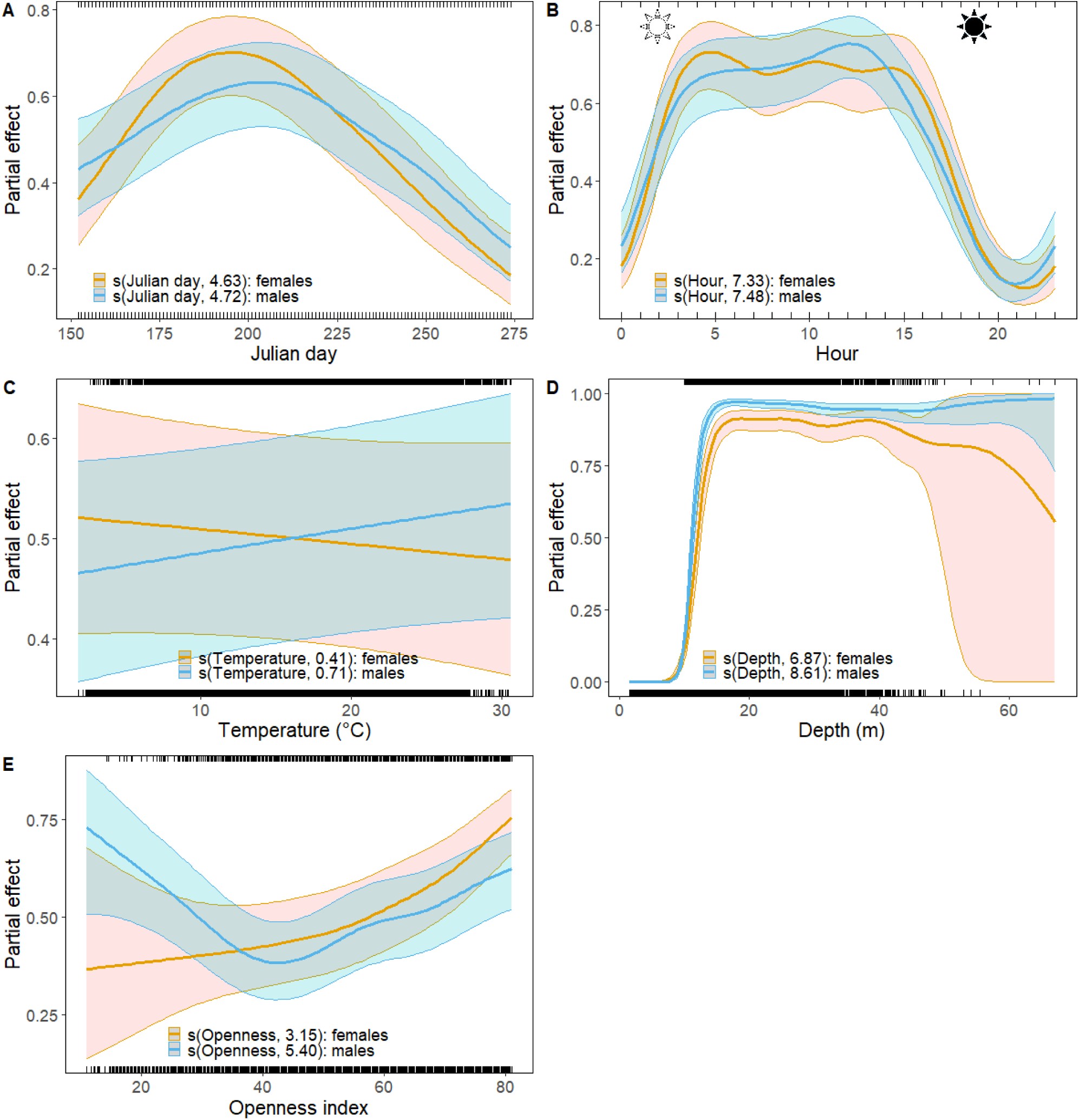
Partial effect plots from the GAMM fitted to the dive data of Saimaa ringed seals in summer. Estimated relationships between the probability of occurrence of foraging behaviour and A) Julian day, B) hour of the day, C) air temperature, D) water depth, and E) openness of the water area (openness index). Females are shown in gold and males in light blue, and the solid line represents the mean and shaded area the 95% confidence intervals. Rugs on the top and bottom axes depict model predicted foraging and non-foraging, respectively. The estimated degrees of freedom for the corresponding model covariate are given in brackets. Sun symbols in figure B) mark average sunrise and sunset times (Coordinated Universal Time, UTC) in the area over June-September.

According to the winter model, males were more likely to forage towards the end of winter (late March) while female foraging probability had a less obvious seasonal trend, with peaks occurring both at the start (early December) and end of the winter data collection period (Fig. 6A). Similar to the summer model, foraging probability peaked in the morning; however, the time window of this higher probability was much narrower compared to the summer period (Fig. 6B). Air temperature had a small but significant negative correlation with foraging probability in males (χ^2^ = 9.74, p < 0.01, edf = 0.90), whereas the relationship was non-significant in females (χ^2^ = 2.15, p = 0.07) (Fig. 6C). The relationship between water depth and foraging probability was significant for both sexes (p < 0.001) with their likelihood of engaging in foraging increasing at depths >7-10 m (Fig. 6D). In the winter, the sexes differed in their relationship between foraging and openness of the water area, with males being more likely to forage in open water while females seemed to prefer more sheltered areas (Fig. 6E).

**Fig. 6.**
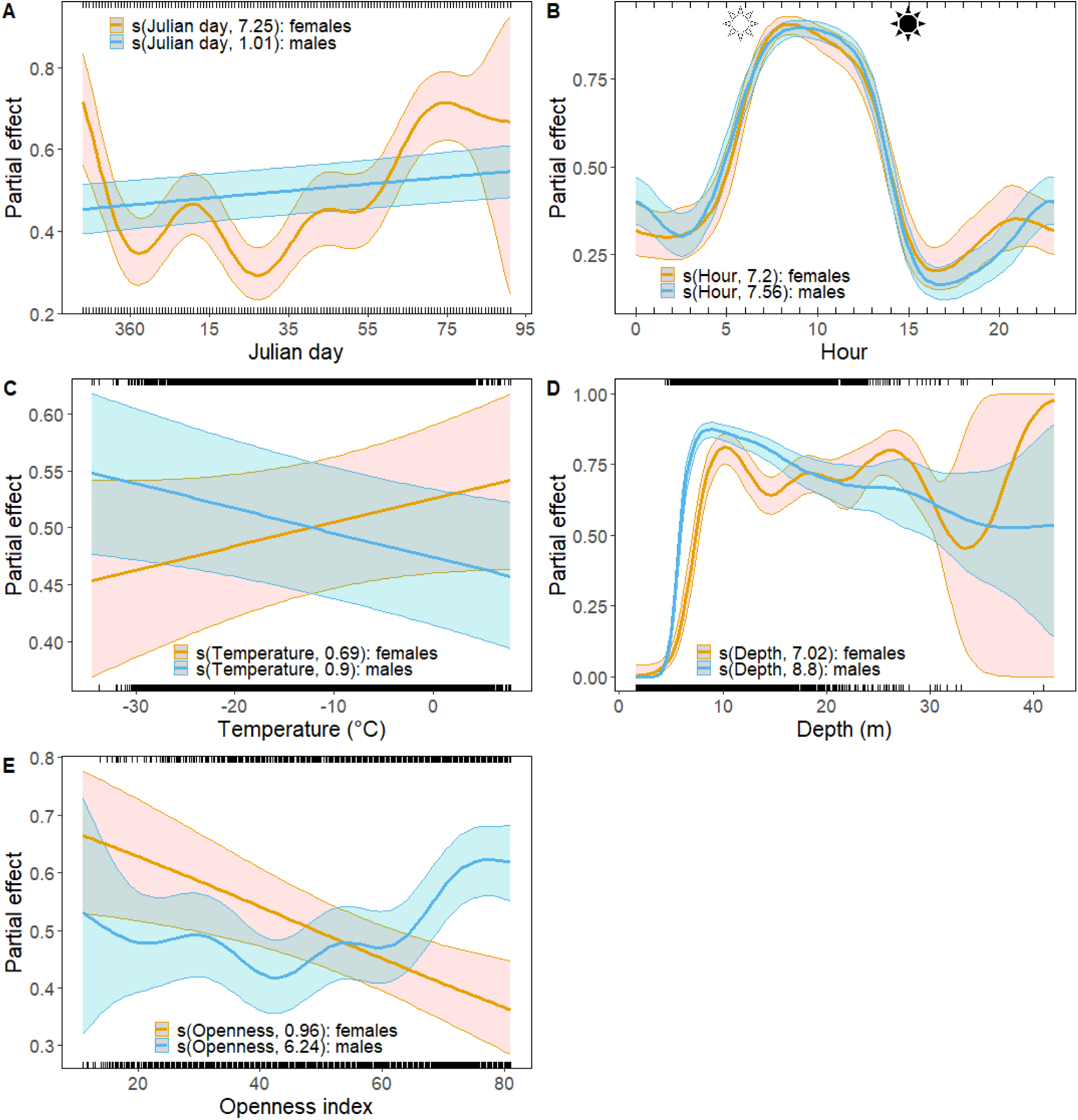
Partial effect plots from the GAMM fitted to the dive data of Saimaa ringed seals in winter. Estimated relationships between the probability of occurrence of foraging behaviour and A) Julian day, B) hour of the day, C) air temperature, D) water depth, and E) openness of the water area (openness index). Females are shown in gold and males in light blue, and the solid line represents the mean and shaded area the 95% confidence intervals. Rugs on the top and bottom axes depict model predicted foraging and non-foraging, respectively. The estimated degrees of freedom for the corresponding model covariate are given in brackets. Sun symbols in figure B) mark average sunrise and sunset times (UTC) in the area over December-March.

Adding the uncertainty resulting from the behavioural state classifications from the HMM did not change the relationship between foraging probability and the covariates in either the summer or winter GAMM and model predictions (Figs. S10 and S11, Supporting information) followed the same pattern as in the partial residual plots (Figs. 5 and 6).

### Goodness-of-fit of the GAMMs

The confusion matrices suggested that both summer and winter GAMMs had a good fit to the data, with an average of 82% correct classifications of foraging and non-foraging states for the summer model and 81% for the winter model. Further, resampling of the HMM states had only a minimal effect on the AUC or the percentage of correct classifications from the confusion matrix: for the summer model, the mean AUC across resamples was 0.885 (0.883−0.887), and the mean percentage of correct classifications of foraging was 87.7% (87.2−88.2%); for the winter model, the mean AUC across resamples was 0.888 (0.886−0.891), while the mean percentage of correct classifications of foraging was 84.2% (83.6−84.6%). Both models performed slightly worse in predicting non-foraging behaviour with the mean percentage of correct classifications of 75.5% (74.9−75.9%) from the summer model and 77.8% (77.3%−78.3%) from the winter model.

### Overlap of foraging areas

Pairwise overlap in the foraging areas (i.e. Bhattacharyya’s affinity) varied from ~0 to 0.912 in the summer, with three pairs of seals, including one female-female pair and two male-male pairs, out of 36 possible pairs sharing more than 25% of the foraging areas. In the winter, only one pair (male-male) shared 61% of the foraging areas, and the BA index was <0.01 for the remaining 14 pairs. When comparing the use of foraging areas within individuals between seasons, the seals used the same but reduced foraging areas in the winter as in the summer, with the mean BA of 0.521 (0.310 - 0.820) among individuals.

## Discussion

### Drivers of foraging

Foraging probability of Saimaa ringed seals peaked in mid-July whereas the winter data showed no clear temporal trend. Harkonen et al. (68) also found a peak in diving activity of Baltic ringed seals in June-July, as opposed to fall and winter when a larger proportion of time was spent hauled out. Although foraging behaviour of ringed seals has not been classified before in such detail as in this study, it has been suggested that they forage mostly during the day based on the fact that Baltic ringed seals were found to spend more time in deeper waters during the day (68). Moreover, von Duyke et al. (69) found repetitive diving, hypothesized to be indicative of foraging, to occur more frequently in the middle of the day under brightest ambient light conditions. The results of the present study corroborate these hypotheses with the highest foraging probability of Saimaa ringed seals occurring during daylight hours in both summer and winter months. In the summer, however, the peak in foraging probability started earlier in the morning and lasted longer through the afternoon compared to winter. During the subarctic mid-summer, the days are long, with sunrise before 5 am, so there is plenty of light in the upper part of the water column for visual predators. Even with the rapid attenuation of light that occurs with increasing water depth in boreal lakes (70,71), pinniped vision is well adapted to low-light levels in deep or murky waters (72). The fact that Saimaa ringed seals concentrated their foraging efforts in the brightest part of the day both in the summer and winter suggests that visual hunting tactics may be important for their foraging success during both seasons. However, the visibility underwater in some parts of the Lake Saimaa can be as little as a few meters, and seals rely also on well-developed mystacial vibrissae while foraging and for orientation (73–75), as there is evidence of blind seals surviving in the lake (75). A key driver for the prolonged foraging in summer compared to winter may be related to replenishing energy stores that are lost during winter (25) and the peak of annual moult in May (during which time they are predominantly hauled out on land) (76,77) while the longer summer days offer an opportunity to maximise foraging efforts through increased visual hunting tactics. In addition, females may also have to replenish the lipid reserves and body mass invested in nursing their pups in spring (78). The female foraging peak occurred a week earlier compared to males, but otherwise there were no temporal differences in the foraging probability between the sexes. Another mechanism underpinning the prolonged foraging during the summer may be linked to movements of prey. Saimaa ringed seals forage on small schooling fish such as roach (*Rutilus rutilus*), perch (*Perca fluviatilis*), vendace (*Coregonus albula*) and smelt (*Osmerus eperlanus*) (24,25). Prey diel activity patterns, such as changes in schooling behavior, vertical movements, and distribution (79–81), may have an effect on the seals’ foraging patterns. Indeed, schooling fish (smelt, vendace, whitefish [*C. lavaretus*] and perch) continue their diel vertical migration in the water column even under ice as a response to prevailing light conditions, and fish were found at the shallowest depths at sunset and sunrise (80), which corresponds to the later start of higher foraging probability of seals in the winter (Fig. 5).

Foraging probability of Saimaa ringed seals was greatest at depths of 10-30 m in the winter and at depths of 15 m and above in the summer. The pelagic fish of Lake Saimaa consists mostly of smelt and vendace (82), which are among the main prey species of the Saimaa ringed seal. Jurvelius et al. (79) found most of the larger smelt (>9 cm) in depths of >25 m in Lake Saimaa, whereas the smaller fish tended to concentrate in the shallower depth layer of 0-17 m. Further, the largest catches of smelt and vendace in the lake were in depth layers of 10-15 m and 25-30 m, respectively (83). It is thus possible that the higher energetic content of these larger fish, coupled with their occurrence in large numbers, is enough to encourage Saimaa ringed seals to dive to deeper depths, particularly in the summer months when vendace are especially lipid rich (84). Moreover, vendace aggregated in deeper water during the daylight hours compared to nighttime (80). Focusing foraging effort to specific depths would be beneficial if it resulted in repeated capture and consumption of aggregated prey, thus maximizing energetic profitability. The dive depths of numerous species of marine mammals, such as Weddell seals, *Leptonychotes weddellii* (85), crabeater seals, *Lobodon carcinophaga* (86), and sperm whales, *Physeter macrocephalus* (87), have been linked to the vertical migration of their pelagic prey.

### Sex-specific foraging strategies

In addition to dynamic environmental conditions, foraging strategies of animals are often influenced by intrinsic factors, such as reproductive state or sex (88–91). Sex-specific foraging behaviour is common in vertebrates, including ungulates (92), pinnipeds (93,94), birds (90,95) and cetaceans (96), and may often be explained by differences in body size between sexes (93,95–97). Unlike grey seals (98) and some arctic ringed seals (99), adult Saimaa ringed seals are not strongly sexually dimorphic in terms of their body length, maximum girth or body mass (100). We found subtle differences between female and male Saimaa ringed seals in some aspects of their foraging behaviour. For example, females had, in general, a slightly higher predicted probability of foraging over the winter compared to males, with a significant difference in early December (Fig. S11, Supporting information). Considering that we only used dives to predict behavioural states (excluding haul-out and surface time periods), it could be speculated that increased foraging probability and time spent foraging are associated with the breeding status of the females and the need to replenish the energy transferred to a pup. Adult grey seal (*Halichoerus grypus*) females in the northwest Atlantic also spent more time foraging (88), whereas female pups had a higher probability of remaining in a foraging state than males in Wales but not in northeast Scotland, suggesting that sex specific behaviour may be modulated by habitat (28). During winter, female Saimaa ringed seals had a higher foraging probability in more sheltered water areas enclosed by islands or land whereas males seemed to prefer open areas. In contrast, female foraging probability increased in the summer with the openness of the water area. This could indicate the existence of sex-specific foraging strategies that are adapted seasonally, for example, to the distribution or occurrence of prey. For example, the limnetic form of European smelt spawns in shallow waters in winter-early spring (101), and this brings large aggregations to sheltered shoreline habitats.

Male Baltic ringed seals were found to dive deeper and over a longer duration compared to females in some areas of the Baltic Sea (68), and the authors hypothesize that this was related to the size of the animals. However, we found no effect of sex on the depth of foraging dives between females and males in Lake Saimaa. Maximum dive depth of ringed seals correlates positively with body mass (102), and the lack of strong sexual dimorphism in Saimaa ringed seals may at least in part explain the contrasting results of this study. Another explanation is that the depth of Lake Saimaa is too shallow for any sex-related physiological differences in dive depth limit to become apparent. Also other factors, such as the seasonal availability of prey or the reproductive state of the animal, may influence the diving depth of air-breathing aquatic animals. For example, mature male ringed seals in the arctic dove to shallower depths and for shorter durations than breeding females or subadult males during the summer breeding season (103), whereas no sex-related differences in dive depth were found during autumn or winter (104).

### Foraging area overlap

Most of the seals that were tagged in this study had largely non-overlapping foraging areas, especially during winter. Niemi et al. (22) also found that the winter core areas (where most of the seals’ activities occur) of adult females did not overlap, providing evidence of competitive exclusion, avoidance, or territoriality over the breeding season. In addition to territoriality, low levels of overlap in the foraging areas may also result from spatial partitioning of limited resources among individuals. Several previous studies have found evidence of demographic habitat or resource partitioning in ringed seals, whereby mature individuals were associated with good ice conditions and juveniles were found near the ice edge or in the open-water areas (69,105,106). Some evidence of demographic habitat partitioning has also been found in Lake Saimaa, where two sub-adult males inhabited the edges of the breeding areas (22). In this study, two adult male seals shared over 90% of the same foraging area in the summer, but the extent of overlap was reduced to about 60% in the winter. It thus seems that the seals may exhibit some degree of resource or habitat partitioning, especially during the winter ice-covered period, when resources may be more limited, or the foraging window may be reduced due to fewer hours of daylight. Similar to this study and a study by Niemi et al. (22), the home ranges of ringed seals in Beaufort and Chukchi Seas were much smaller (most of them within 3 km^2^) in winter (107), compared to the summer, when seals were performing extensive foraging trips. However, unlike the seals in this study, arctic ringed seals showed extensive overlap in their home ranges during the subnivean period, with individuals sharing the use of breathing holes (107).

### Time budget and possible consequences of climate change

Overall, Saimaa ringed seals allocated between a fifth (in winter) and a third (in summer) of their diving time to dives that fit the characteristics of foraging behaviour. The reduction in time allocated to foraging in winter is in agreement with evidence from stable isotope analysis that Saimaa ringed seals lose weight during winter and spring (25). Nevertheless, even the higher proportion of time budget allocated to foraging in the summer is much lower than what was estimated for arctic ringed seals, which were hypothesized to forage on average over 12 h/day from August through January (69). This may suggest that prey species in the lake for Saimaa ringed seals are abundant. Moreover, as a generalist predator (25), the Saimaa ringed seal can feed on several different species of fish and switch target prey depending on their availability. The fact that the seals were foraging in both open and sheltered water areas is indicative of flexible foraging strategies, either through diverse foraging tactics that are adapted to the bathymetric and dynamic hydrological features of their environment, or through foraging on a range of species that occur in different habitats and/or may display different anti-predatory behaviours. The proportion of time spent engaged in shallow short dives with little vertical sinuous movements (*i.e.*, the inferred shallow inactive state) was over 45% in the winter compared to just 19% in the summer. These dives occurred in both deep and shallow water and were not tied to any specific behaviour. We suspect that these shallow dives are linked to resting at lair sites and/or maintenance of breathing holes and are ultimately connected to the more restricted habitat use during winter compared to summer.

The deep and long sleeping/resting dives that composed 9% of all dives in the summer were missing entirely in the winter period. In winter, ringed seals dig subnivean lairs for resting, pupping and nursing, and the lack of sleep dives, combined with the fact that the proportion of time that Saimaa ringed seals spend hauled out increases in the winter (Fig. S12, Supporting information), may indicate a preference to rest/sleep in snow-lairs rather than in water. Indeed, the temperature in snow lairs is considerably warmer than the external environment and can reach more than 5°C when occupied by a seal (108). This, together with the fact that sleep in mammals is accompanied by a decrease in body temperature (109) and that heat transfers ~24 times faster in water than in air, would make sleeping in a snow lair energetically more efficient. It may be that Saimaa ringed seals conserve energy by sleeping in snow lairs, as the proportion of time spent in foraging was estimated to be considerably lower during winter compared to summer. It is also possible that sleep occurred in water but at depths less than the dive limit (<1.5 m) that was set by the tag manufacturer, as was observed in Kunnasranta et al. (27). Also the increased proportion of time spent at the surface between dives (but not hauling out) supports this (Fig. S13, Supporting information).

The length of thermal winter, defined as the period when daily mean temperature falls below 0℃, is forecasted to shorten by 30-50 days in the area occupied by Saimaa ringed seals in 2040–2069 under an intermediate climate scenario (RCP4.5) by the Intergovernmental Panel on Climate Change (110). Further, the probability of absence of thermal winter is forecasted to increase from current 0-2% to 5-20%. These changes will undoubtedly have drastic effects on the population of ice-obligate Saimaa ringed seals, mediated by increased pup mortality due to the combined effect of premature collapse of pupping lairs during early onset of spring and increased vulnerability of pups to predation during winters that are completely without snow (16). In fact, a recent study projected median ringed seal population declines ranging from 50% to 99% by the end of the century, accompanied by substantial changes in population structure (111). In addition to population-level effects, climate change will likely have an impact on the behaviour patterns and activity budgets of Saimaa ringed seals, via changes in thermoregulation as the seals cannot conserve energy by resting in snow lairs, or via more indirect changes. For example, we hypothesize that during winters with poor snow conditions, Saimaa ringed seals will have to allocate more time (and energy) into finding suitable nest sites and constructing lairs. Moreover, females will face the issue of nursing their pups without shelter provided by the pupping lair. Climate change may also pose other indirect risks, such as reduced availability of prey species due to shifts occurring in food webs stimulated by warmer water temperatures. For example, elevated temperatures seem to favour warm-water fish species in northern latitudes while cold-water species, such as vendace, tend to suffer (112).

### Conservation implications

Saimaa ringed seals are thought to be most sensitive to disturbance during the breeding season (113). The higher proportion of time spent hauled-out in winter shown in this study highlights the significance of sufficient snow and ice cover for subnivean structures used for resting and pupping. However, when snow conditions are poor during mild winters, the seals are forced to rest in water or on open ice, which is not only energetically more demanding, but also exposes them to increased disturbance and predation. Future management plans should thus keep focusing on minimizing human disturbance and predation pressure during the sensitive winter season in key habitats. Incidental bycatch mortality is still a major threat for the Saimaa ringed seal population, and this study provides new knowledge supporting the need for implementing broader fishing restrictions. Our results indicate that Saimaa ringed seals’ foraging activity peaks in July, thus coinciding with the end of the springtime fishing closure, a 2.5 month long period during which fishing with gillnets is banned in Lake Saimaa. Extending the fishing closure has been proposed on the basis of elevated bycatch mortality of juveniles in July (18). The present study provides further evidence of intense diving activity by the seals in late summer, which increases the risk of entanglement in fishing gear. In addition, we show that the seals use the entire water column and various habitats of the lake for foraging, *i.e*., that there are no ‘seal-safe’ areas for gillnet fishing. It is also notable that seals forage mostly during daylight, which contradicts some assumptions on ‘seal-safe’ fishing by using gillnets only during daytime.

## Conclusions

In this study, we demonstrate for the first time, the profound influence of environmental variation on the behavioural strategies of ringed seals. Using a combination of HMMs and GAMMs applied to telemetry data collected on individual dives, we show contrasting activity budgets in summer versus winter. We reveal that the Saimaa ringed seals’ primary activity (36% of time) in summer months is foraging, whereas this is the least dominant activity in winter (21% of time). Moreover, aquatic resting behaviour, in the form of long and relatively deep dives, was present in summer but not in winter, demonstrating behavioural plasticity in resting strategies in relation to environmental conditions. We suggest that foraging behaviour of Saimaa ringed seals is largely influenced by diel vertical movements and availability of fish, and that the seals optimize their energy acquisition while conserving energy, especially during the cold winter months.

Future studies should investigate the behavioural patterns of juvenile seals, which is the demographic portion of the population that is most vulnerable to bycatch mortality. Ultimately, predictive modeling that integrates knowledge of the demography, genetics and ecological plasticity of Saimaa ringed seals in response to environmental variation will be required to assess how climate change will affect this endemic subspecies.

## Acknowledgements

We are grateful to Tuomas Rajala for the custom R-script to retrieve weather variables. Milaja Nykänen was funded by the Finnish Cultural Foundation. Matt Carter was supported by the INSITE EcoSTAR project (Natural Environment Research Council; NE/T010614/1). Data collection was funded by Saimaa seal LIFE (LIFE12NAT/FI/000367), Raija and Ossi Tuuliainen Foundation (#2485/02.07.02/2010) and WWF Finland (#2421/02.07.02/2010).

## Ethics approval

The seals were captured and handled for the deployment in accordance with the Finnish environmental authorities, Centre for Economic Development, Transport and the Environment (ESAELY/433/07.01/2012 and ESA-2008-L-519-254), and the Project Authorisation Board (ESAVI/8269/04.10.07/2013 and ESAVI-2010-08380/Ym-23).

