## Supporting information for "Linking ringed seal foraging behaviour to environmental variability"

Table S1. Summary of GPS-GSM data used to run summer and winter models for the Saimaa ringed seals.

| Seal ID | Sex | Weight (kg) | Summer (open water) | | | | | Winter (ice-cover) | | | | |
|---|---|---|---|---|---|---|---|---|---|---|---|---|
| | | | Start date | End date | Duration (days) | Mean # dives/day | Mean # locations/ day | Start date | End date | Duration (days) | Mean # dives/day | Mean # locations/ day |
| TO-07 | M | 55 | 2008-06-03 | 2009-09-30 | 120 | 255 | 15 | 2009-12-14 | 2010-03-29 | 105 | 239 | 3 |
| VI-09 | M | 124 | 2008-06-01 | 2009-09-30 | 122 | 150 | 23 | - | - | - | - | - |
| OL-10 | F | 59 | 2010-06-01 | 2010-09-30 | 122 | 174 | 12 | 2010-12-01 | 2011-04-04 | 124 | 116 | 7 |
| ER-11 | M | 66 | 2011-06-01 | 2011-09-22 | 113 | 189 | 29 | - | - | - | - | - |
| TE-07 | F | 52 | 2011-06-01 | 2011-09-30 | 121 | 146 | 28 | 2012-01-01 | 2012-02-10 | 40 | 221 | 12 |
| AS-12 | M | 57 | 2012-06-01 | 2012-09-30 | 122 | 240 | 25 | 2012-12-01 | 2013-01-11 | 41 | 291 | 4 |
| ER-09 | F | 42 | 2012-06-02 | 2012-09-30 | 121 | 181 | 23 | - | - | - | - | - |
| VO-12 | M | 63 | 2012-06-01 | 2012-09-30 | 121 | 305 | 10 | 2012-12-01 | 2013-04-14 | 135 | 212 | 2 |
| MI-13 | M | 57 | 2013-06-03 | 2013-09-30 | 119 | 227 | 8 | 2013-12-01 | 2014-03-31 | 121 | 360 | 4 |

Table S2. Summary of candidate HMMs.

| Season | Model | Covariates | AIC | ΔAIC |
|---|---|---|---|---|
| Summer (open water) | m4 | ~bs(Julian day, df = 6) + bs(hour, df = 6) + depth + openness | 986422.4 | 0 |
| | m5 | ~bs(Julian day, df = 6) + bs(hour, df = 6) + temperature + depth + openness | 986433.1 | 10.7 |
| | m3 | ~bs(Julian day, df = 6) + bs(hour, df = 6) + openness | 994165.3 | 7742.9 |
| | m2 | ~bs(Julian day, df = 6) + bs(hour, df = 6) | 995307.4 | 8885 |
| | m1 | ~bs(Julian day, df = 6) | 997794 | 11371.6 |
| | m0 | NA | 998661.6 | 12239.2 |
| Winter (ice-cover) | m5 | ~bs(Julian day, df = 6) + bs(hour, df = 6) + temperature + depth + openness | 433353.2 | 0 |
| | m4 | ~bs(Julian day, df = 6) + bs(hour, df = 6) + depth + openness | 433381.4 | 28.2 |
| | m3 | ~bs(Julian day, df = 6) + bs(hour, df = 6) + openness | 434221.6 | 868.4 |
| | m2 | ~bs(Julian day, df = 6) + bs(hour, df = 6) | 434589.2 | 1236 |
| | m1 | ~bs(Julian day, df = 6) | 436592.4 | 3239.2 |
| | m0 | NA | 436876 | 3522.8 |

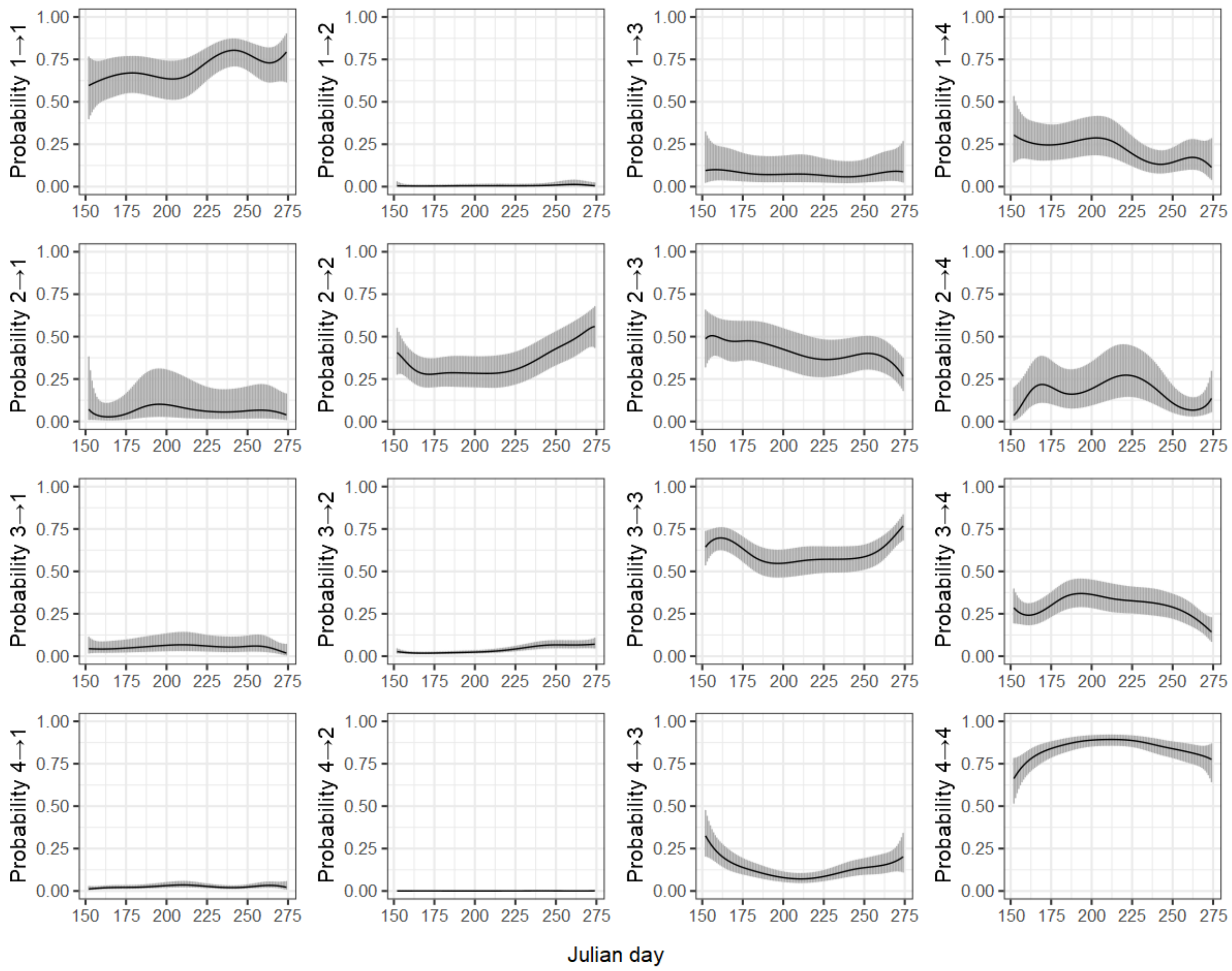


Fig. S1. Plots of the transition probabilities between behavioural states from the summer HMM as a function of Julian day. State 1 = rest/sleep, state 2 = shallow inactive, state 3 = transit/other, state 4 = foraging. The plots show the average trend with a 95% confidence interval.

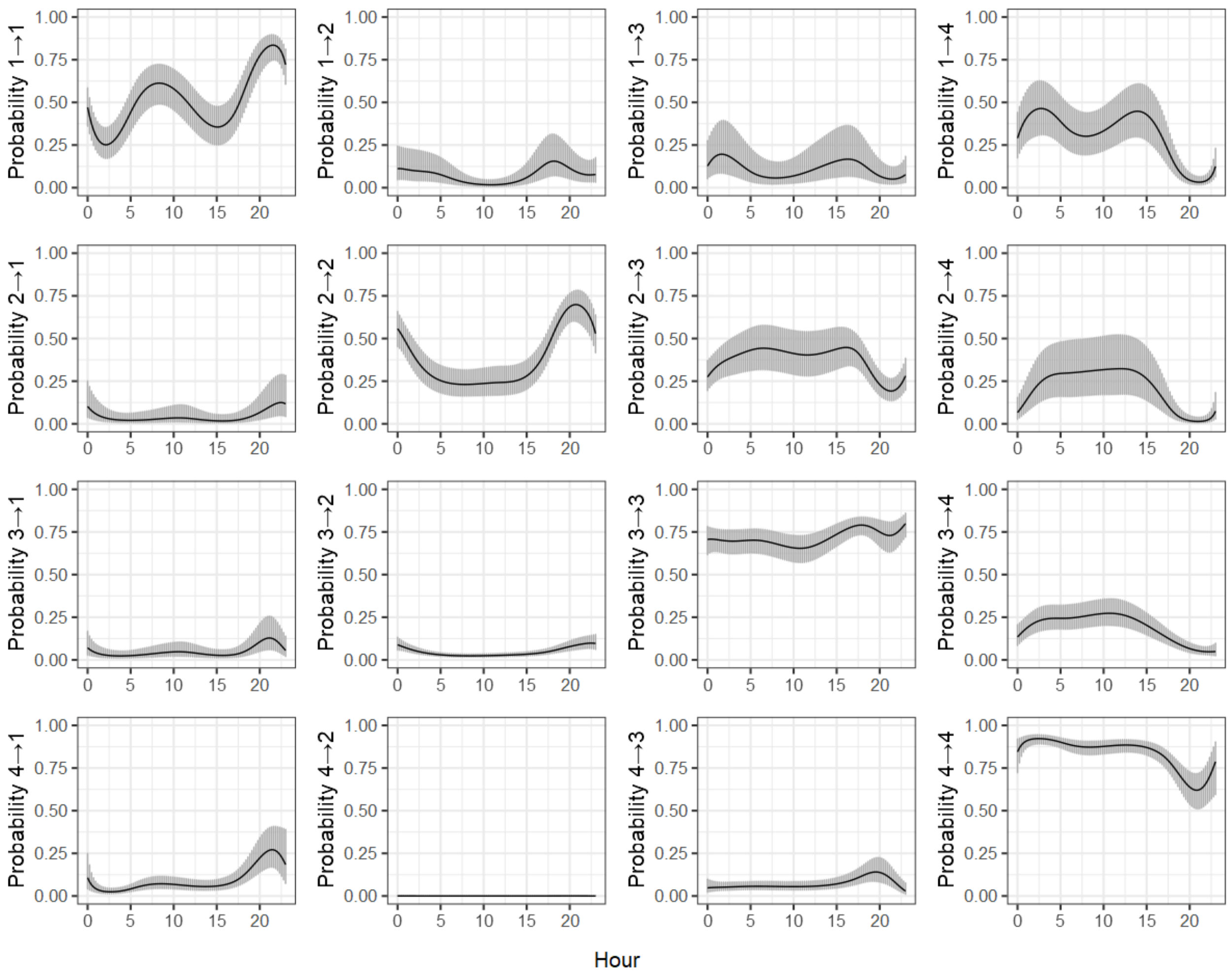


Fig. S2. Plots of the transition probabilities between behavioural states from the summer HMM as a function of hour of the day. State 1 = rest/sleep, state 2 = shallow inactive, state 3 = transit/other, state 4 = foraging. The plots show the average trend with a 95% confidence interval.

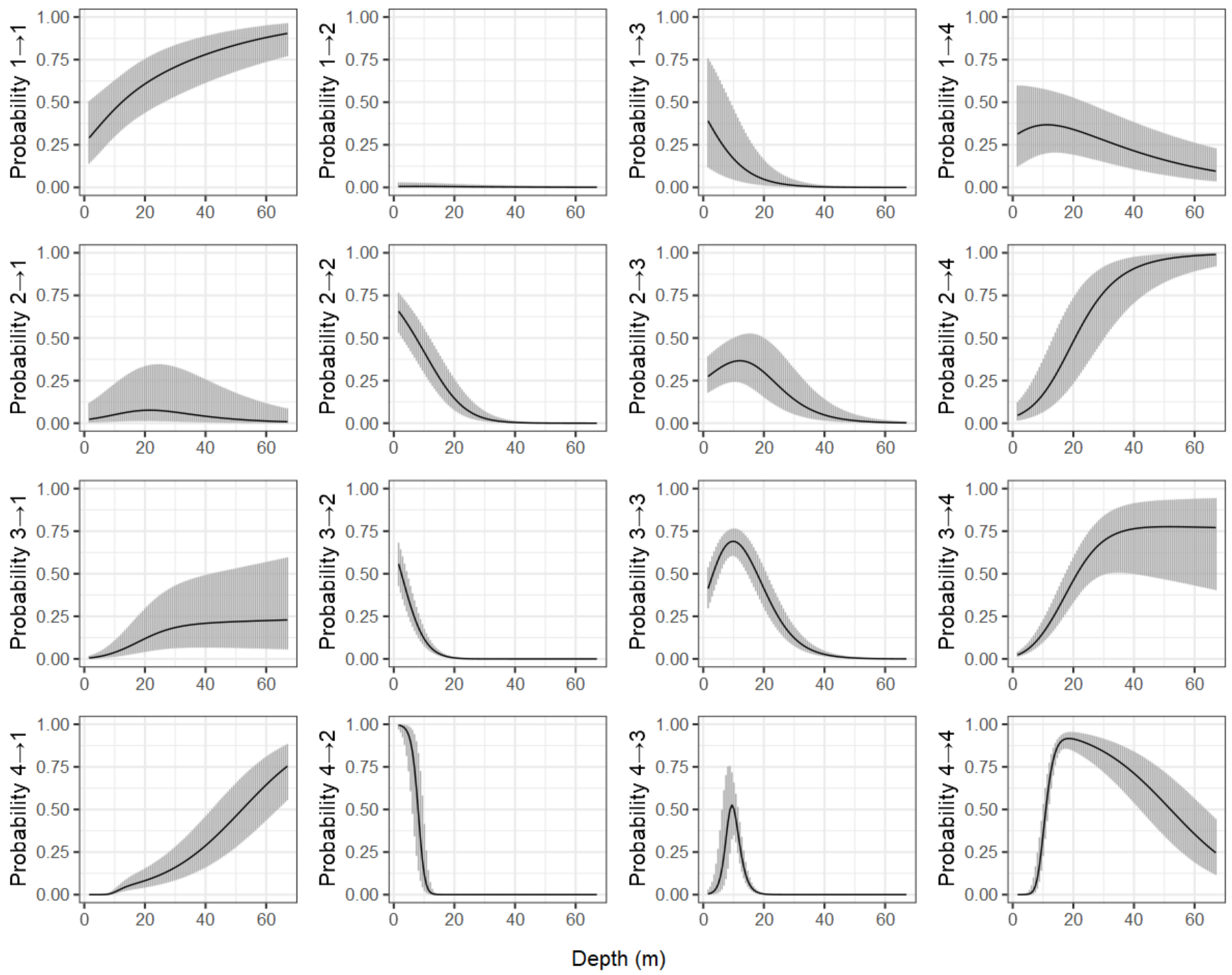


Fig. S3. Plots of the transition probabilities between behavioural states from the summer HMM as a function of water depth. State 1 = rest/sleep, state 2 = shallow inactive, state 3 = transit/other, state 4 = foraging. The plots show the average trend with a 95% confidence interval.

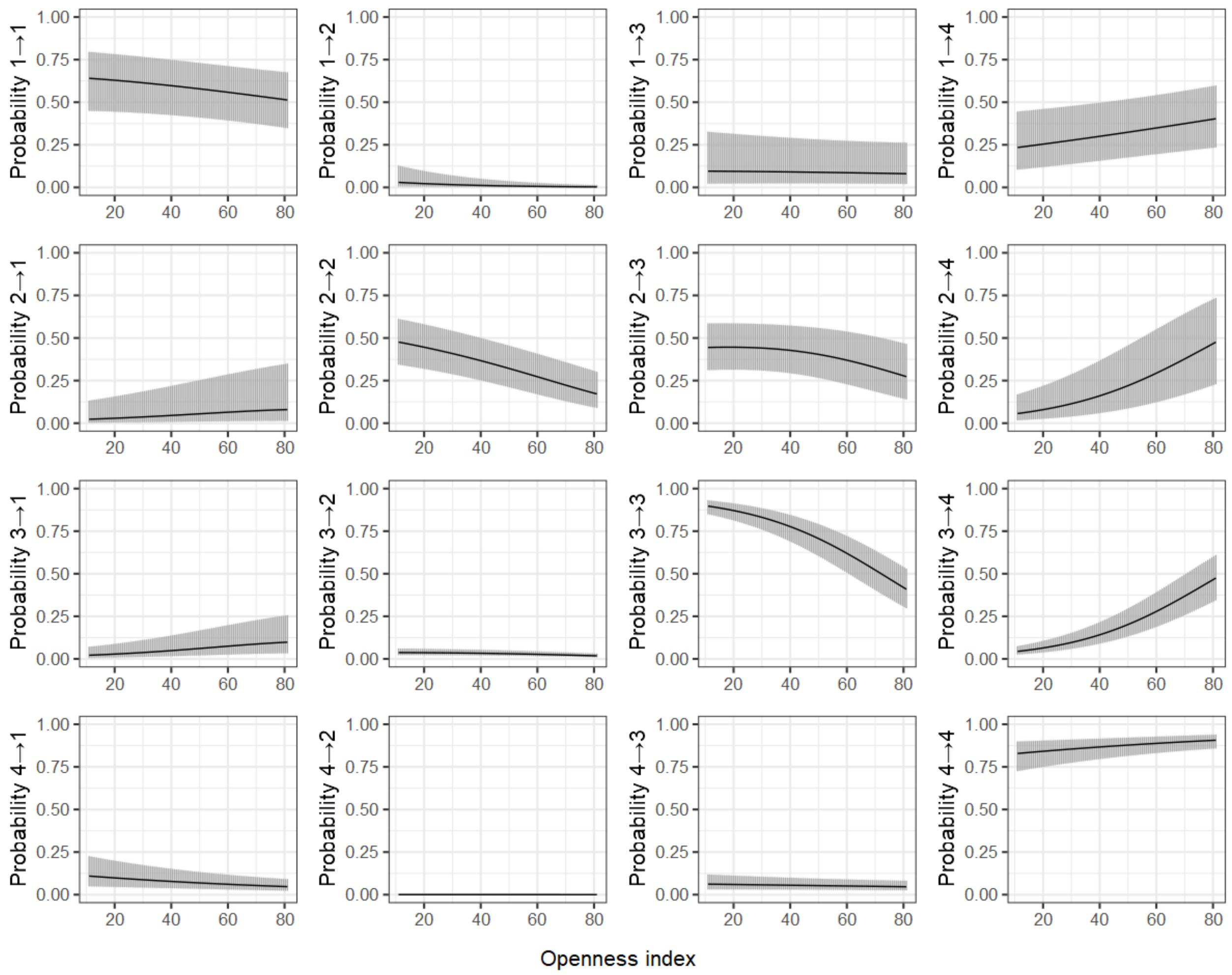


Fig. S4. Plots of the transition probabilities between behavioural states from the summer HMM as a function of openness of the water area (openness index). State 1 = rest/sleep, state 2 = shallow inactive, state 3 = transit/other, state 4 = foraging. The plots show the average trend with a 95% confidence interval.

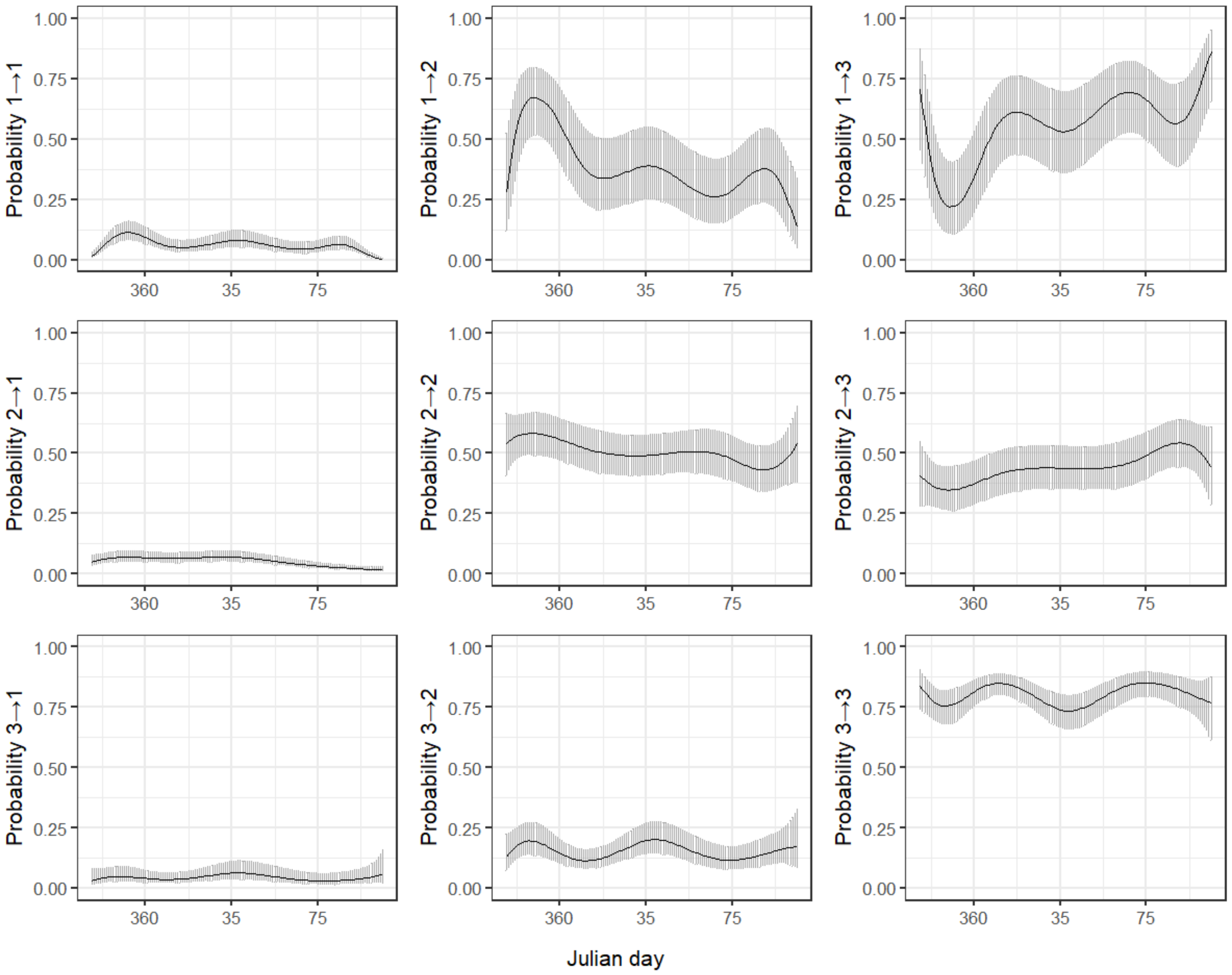


Fig. S5. Plots of the transition probabilities between behavioural states from the winter HMM as a function of Julian day. State 1 = shallow inactive, state 2 = transit/other, state 3 = foraging. The plots show the average trend with a 95% confidence interval.

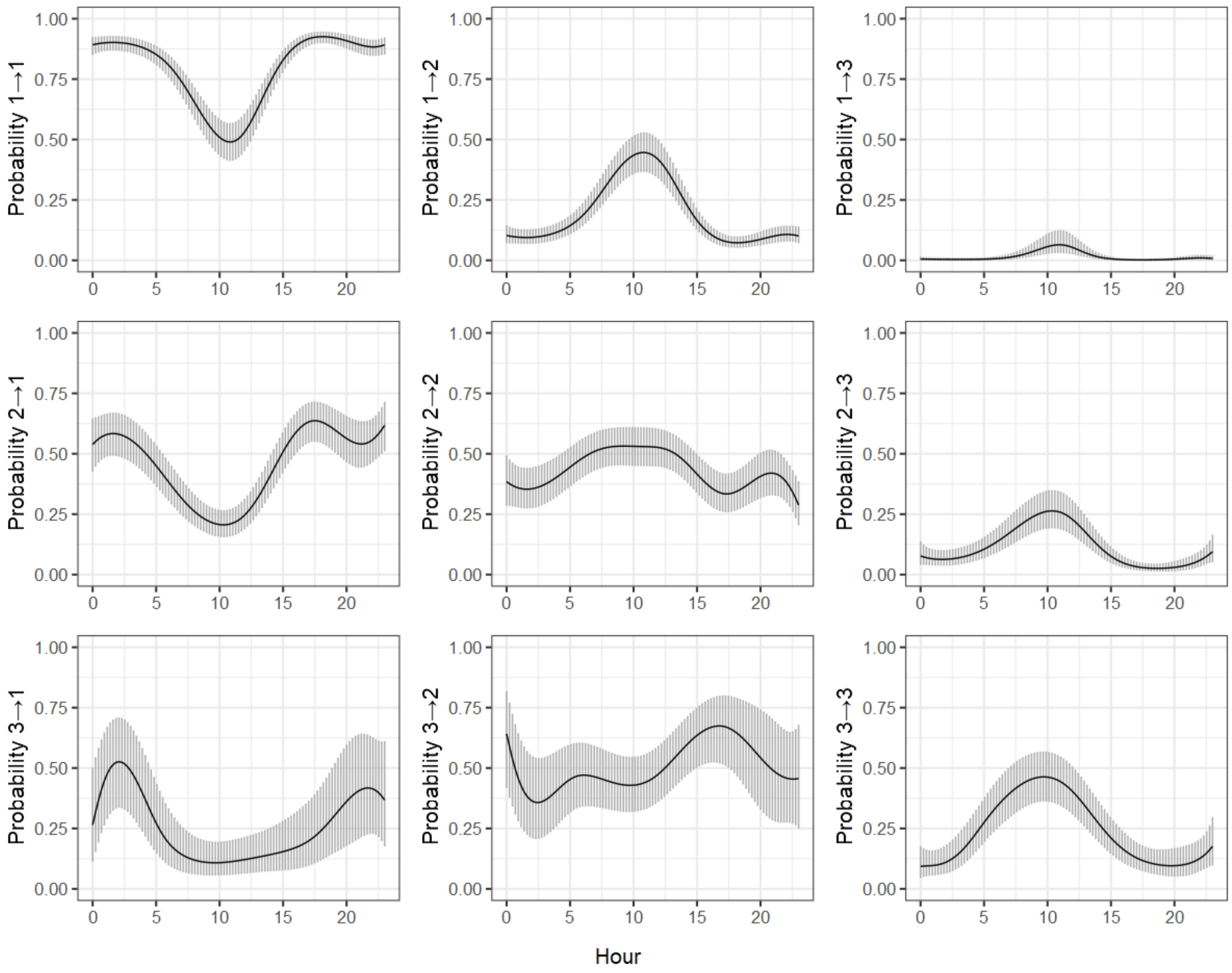


Fig. S6. Plots of the transition probabilities between behavioural states from the winter HMM as a function of hour of the day. State 1 = shallow inactive, state 2 = transit/other, state 3 = foraging. The plots show the average trend with a 95% confidence interval.

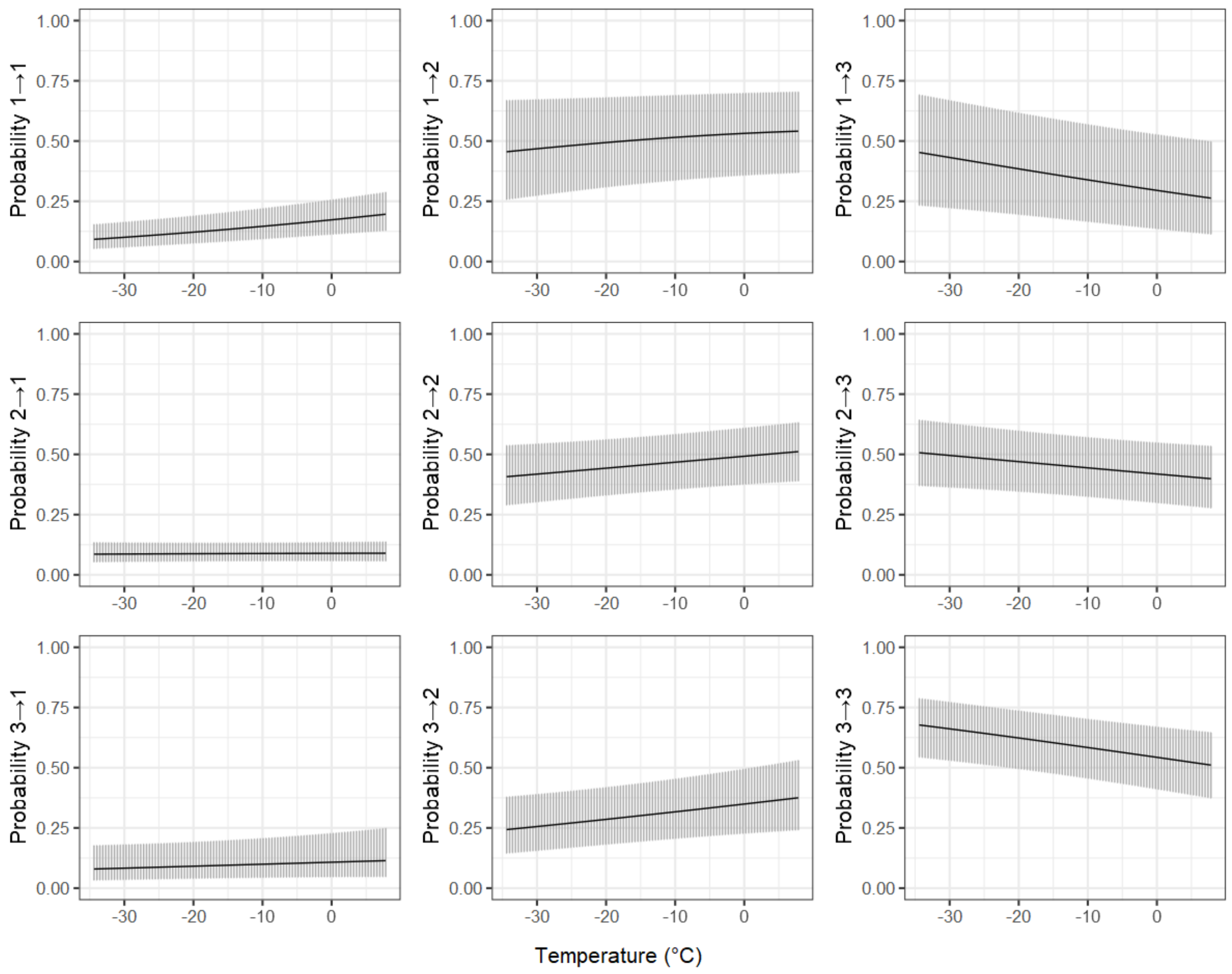


Fig. S7. Plots of the transition probabilities between behavioural states from the winter HMM as a function of air temperature. State 1 = shallow inactive, state 2 = transit/other, state 3 = foraging. The plots show the average trend with a 95% confidence interval.

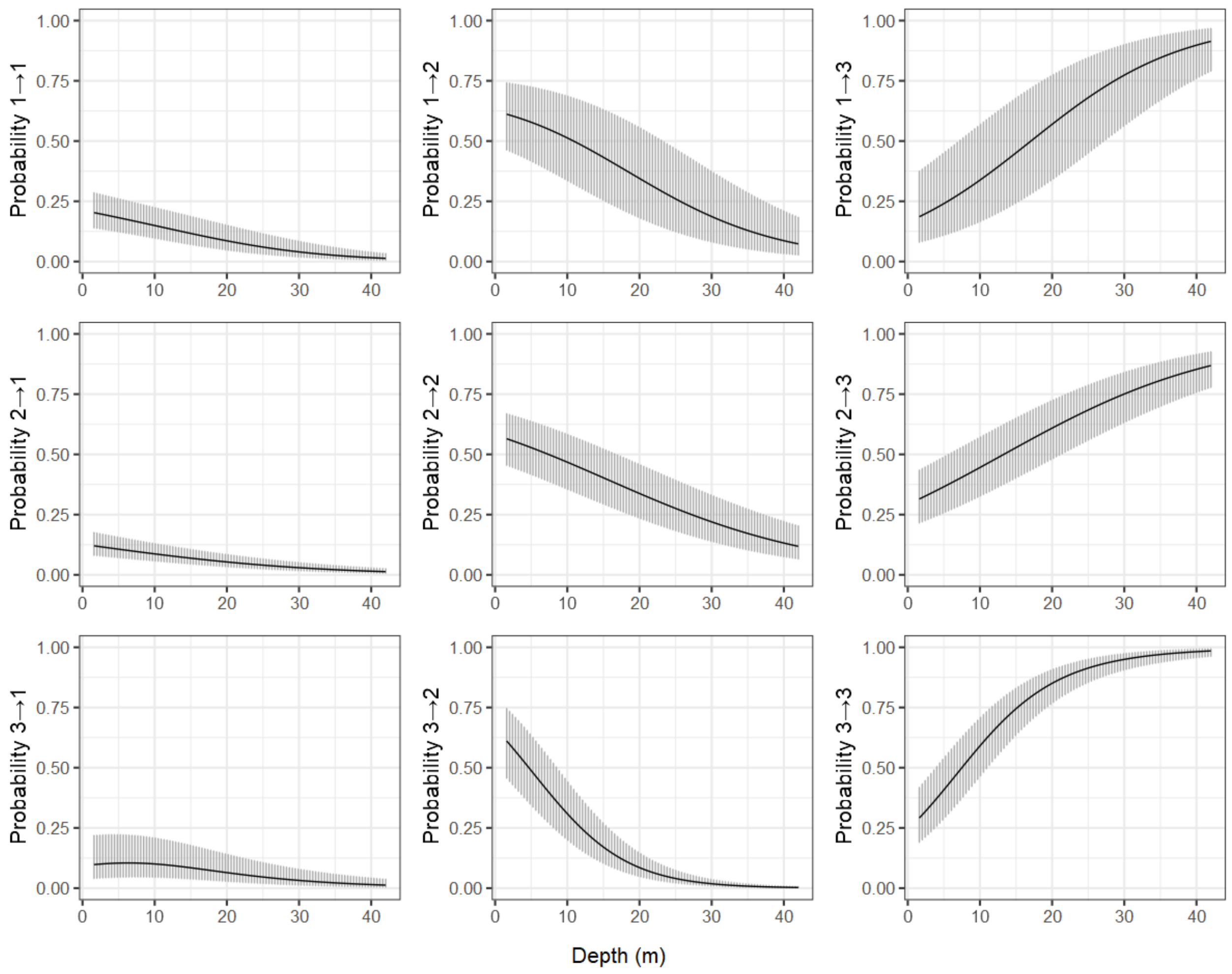


Fig. S8. Plots of the transition probabilities between behavioural states from the winter HMM as a function of water depth. State 1 = shallow inactive, state 2 = transit/other, state 3 = foraging. The plots show the average trend with a 95% confidence interval.

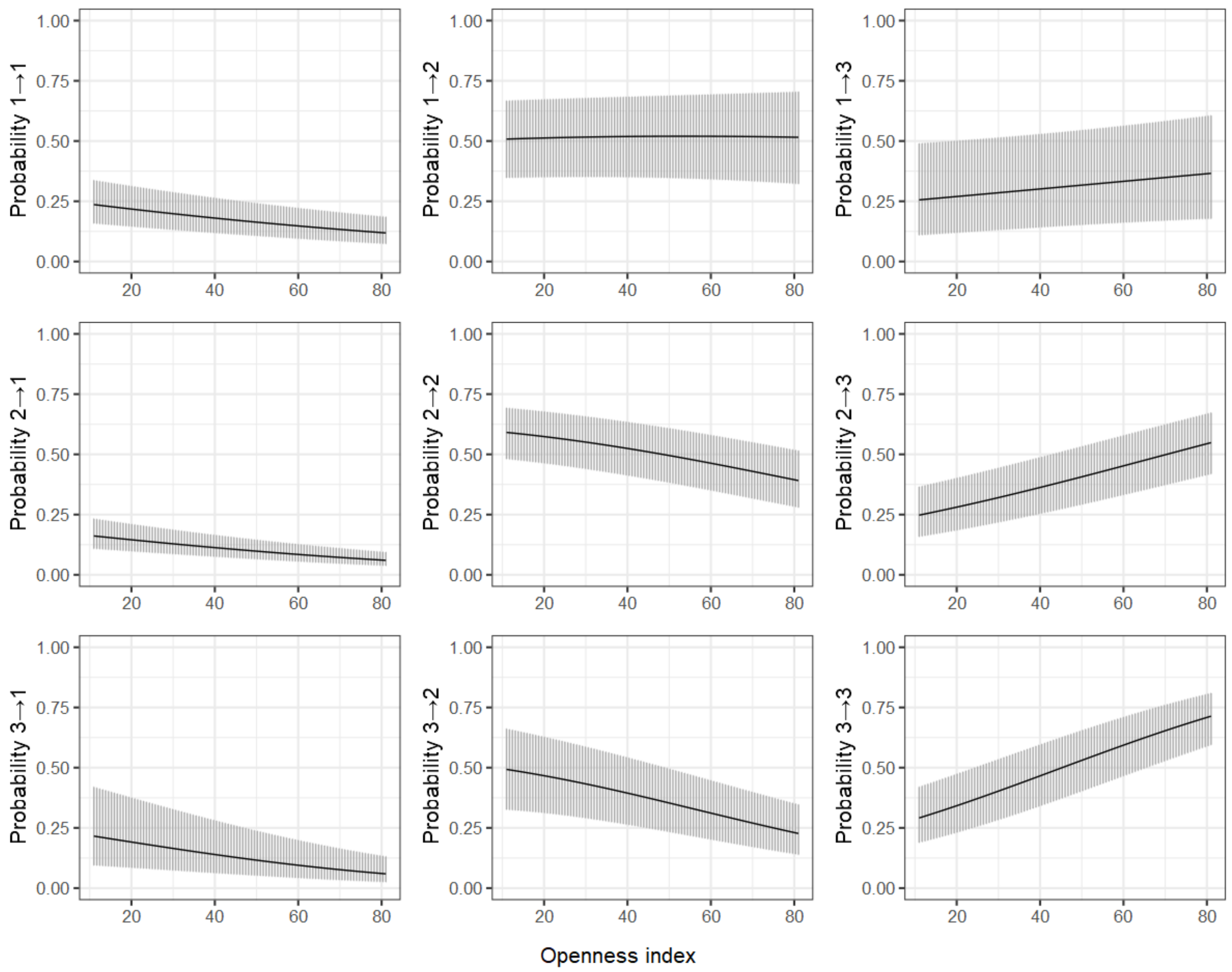


Fig. S9. Plots of the transition probabilities between behavioural states from the winter HMM as a function of openness of the water area (openness index). State 1 = shallow inactive, state 2 = transit/other, state 3 = foraging. The plots show the average trend with a 95% confidence interval.

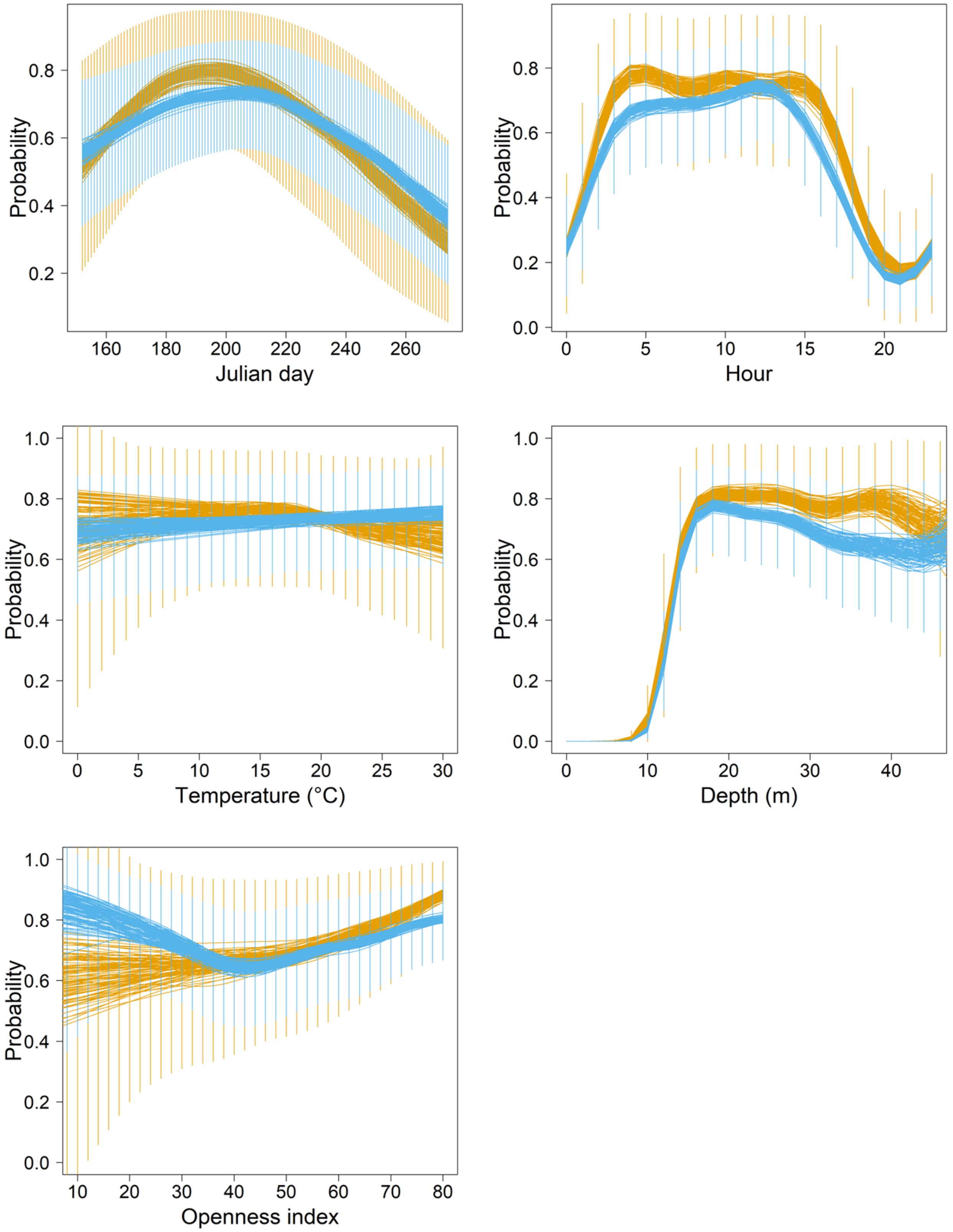


Fig. S10. Predicted probability (mean with 95% prediction interval) of occurrence of foraging behaviour based on the final model for summer for females (gold) and males (light blue) after 100 repeated runs. Predictions were made for each model covariate by holding other covariates at their mean.

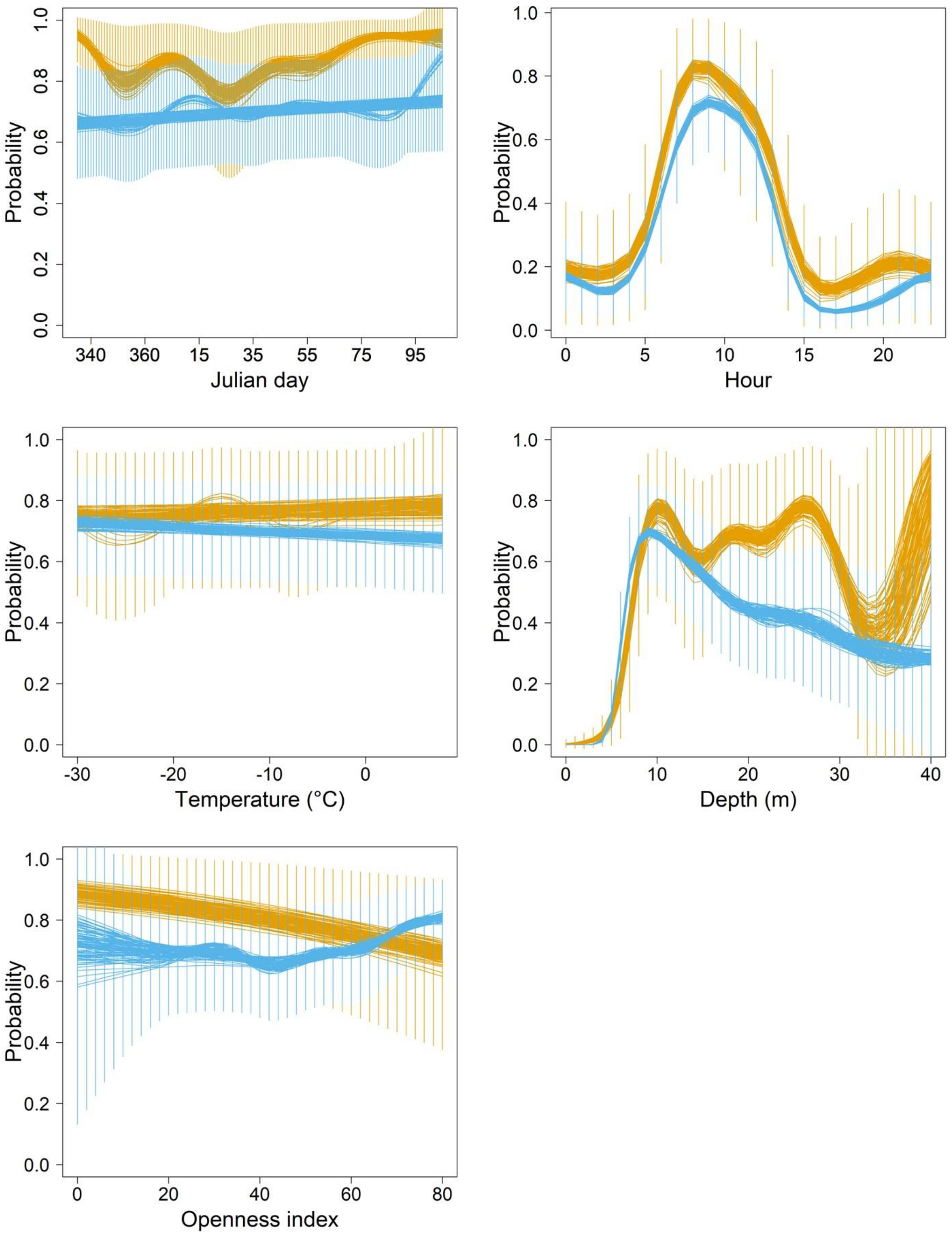


Fig. S11. Predicted probability (mean with 95% prediction interval) of occurrence of foraging behaviour based on the final model for winter for females (gold) and males (light blue) after 100 repeated runs. Predictions were made for each model covariate by holding other covariates at their mean.

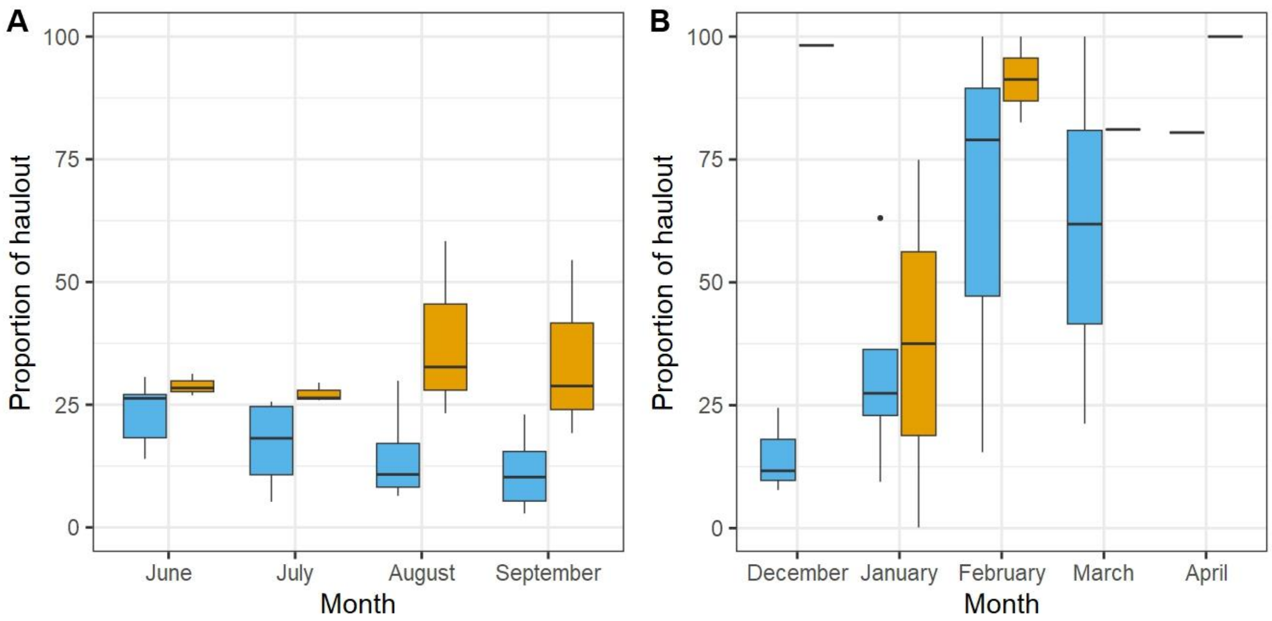


Fig. S12. The proportion of time spent hauled out during a) summer and b) winter seasons. Males are shown in light blue and females in gold colour.

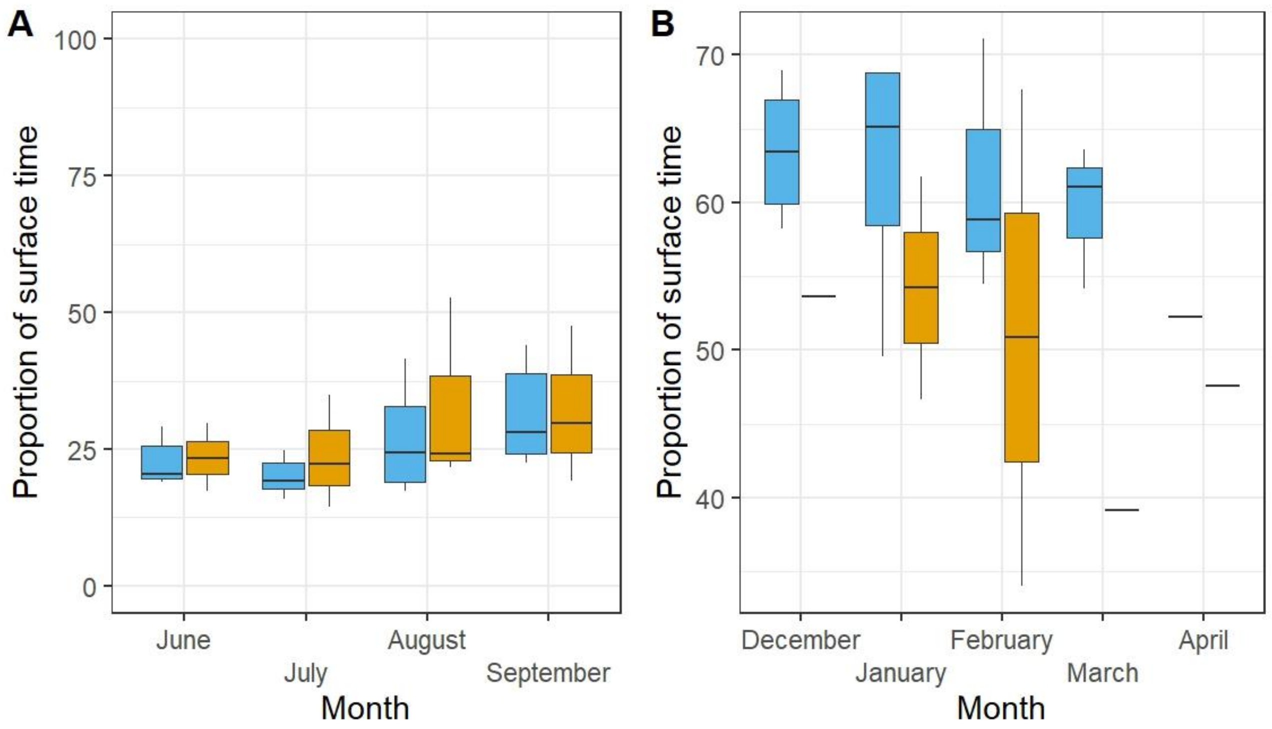


Fig. S13. The proportion of time spent on the surface between dives (but not hauled out) during a) summer and b) winter seasons. Males are shown in light blue and females in gold colour.
